# Phenotypic divergence is driven by mobile genetic elements in a heritable insect symbiont

**DOI:** 10.64898/2025.12.19.695180

**Authors:** Balig Panossian, Matthew R. Kolp, Taoping Wu, Keertana Tallapragada, Vilas Patel, Elliott B. Goldstein, Kerry M. Oliver, Lee M. Henry, Benjamin J. Parker

## Abstract

Heritable microbes profoundly influence insect biology, yet the traits they confer often evolve rapidly and differ among closely related symbiont strains. Despite their importance, we lack a clear understanding of how novel traits arise in symbiosis and how this diversity influences host ecology in natural populations. The aphid facultative symbiont *Regiella insecticola* is ideally suited to this question because of its strong lineage-specific variation in host benefits. By generating 20 high-quality genomes, we found that *Regiella*’s evolution is driven largely by gene gains mediated by mobile genetic elements. We identified a plasmid (pRILSR1) that encodes a type IV secretion system and a highly expressed predicted effector that has been convergently acquired by *Regiella* strains from pea aphids. Notably, only pRILSR1-bearing strains confer protection against the specialist fungal pathogen *Pandora neoaphidis*, indicating that gains and losses of the plasmid underlie the evolution of this key defensive phenotype. Using a multi-year field study, we further show that the pRILSR1 plasmid is strongly associated with *Regiella* found in pea aphid populations adapted to specific host plants, driving variation in symbiont-mediated defense across populations. Together, our results show that mobile genetic elements generate key adaptive traits in microbial symbionts and, in doing so, drive phenotypic divergence among host populations.

## Introduction

Symbioses between microbes and eukaryotic hosts are ubiquitous [1]. These associations range from open systems shaped by continual colonization to closed systems in which symbionts are strictly vertically transmitted and isolated from the environment [2]. This variation fundamentally shapes symbiont evolution. In open systems like animal guts, mobile genetic elements (MGEs) drive rapid genetic change and shape symbiont phenotypes [3]. In contrast, obligate heritable symbionts evolve under limited gene flow and strong drift, leading to genome reduction and functional decay [4].

Many insect–microbe symbioses fall between these extremes. Vertical transmission promotes gene loss, yet occasional horizontal transfer introduces new genes that fuel functional innovation [5-7]. This pattern is exemplified by heritable facultative symbionts of insects, which are not essential for host survival and tend to occur at intermediate frequencies in natural populations [8]. Key factors shaping the persistence and spread of these microbes are the ecological benefits they provide to insect hosts, such as protection against natural enemies [6]. In facultative symbionts, genes encoding host benefits have been shown to be located on MGEs, including plasmids and bacteriophages [9, 10], and horizontal transfer of MGEs has the potential to drive host functional diversification.

Recent research has shown that symbiont phenotypes evolve rapidly, with closely related strains exerting markedly different effects on their hosts. For example, while *Wolbachia* typically spreads through reproductive manipulation, some *Drosophila*-infecting strains protect against viruses, with the strength of protection varying across symbiont strains [11]. Similar strain-dependent variation has been observed in *Spiroplasma*, where closely related lineages differ in their ability to defend aphids against natural enemies [12], as well as in other insect symbiont systems [13, 14]. These findings suggest that insect symbionts frequently gain and lose genes that underlie insect phenotypes. A key challenge is to determine how ecologically relevant traits arise in symbionts and whether they evolve gradually within lineages or spread rapidly through gene exchange.

Aphids and their symbionts provide a powerful system to address this question. In addition to their obligate symbiont *Buchnera aphidicola*, which supplies essential nutrients missing from plant phloem [15], aphids harbor several facultative symbionts that are primarily vertically transmitted but can also move horizontally between host lineages [16]. *Regiella insecticola* (hereafter *Regiella*) stands out for its diverse, lineage-specific ecological benefits, including protection against fungal pathogens [17, 18], defense against parasitoid wasps [19], resistance to a viral pathogen [20], and improved performance on certain host plants [21]. In pea aphids (*Acyrthosiphon pisum*), *Regiella* is consistently associated with specific plant-adapted aphid populations (called ‘biotypes’) [22-24]. Yet, we still do not understand how *Regiella* acquires and diversifies the genes underlying these benefits, or how this diversity is linked to its hosts’ ecology.

In this study we combine culture-facilitated long- and short-read genome sequencing of diverse *Regiella* strains with laboratory and field assays to understand the gains and losses of a key adaptation— protection against a fungal pathogen. We found that mobile genetic elements have played a critical role in *Regiella*’s evolutionary history. In particular, a small plasmid (called pRILSR1) encoding a complete *virB*-like type IV secretion system (T4SS) and highly expressed predicted effector has been horizontally exchanged among distantly related clades associated with pea aphids. Critically, we demonstrated that the pRILSR1 plasmid is essential for *Regiella*-mediated protection against the fungal pathogen *Pandora neoaphidis* (hereafter, *Pandora*). In results from a multi-year field study, we find that the plasmid is strongly associated with *Regiella* from specific plant-adapted pea aphid populations— enhancing their fungal resistance—indicating the symbiont is driving phenotypic divergence among aphid biotypes. Together, our results suggest that mobile genetic elements play a central role in shaping adaptive trait evolution in facultative symbionts and that this variation is intimately tied to host ecology.

## Results

### *Regiella* is highly genetically diverse as shown by multi-locus sequence type (MLST) analysis

We first used an MLST scheme to characterize the genetic diversity and host associations of *Regiella* across 23 aphid species (Table S1) based on [25]. We identified 119 unique sequence types among 347 *Regiella*-positive aphids. Qualitatively, we find that some *Regiella* strains were restricted to a single aphid species, while others spanned multiple species (Figure S1). Similarly, some aphid species harbored only a single strain of *Regiella* while others carried multiple strains (Figure S2). These patterns suggest that *Regiella* symbionts are gained and lost among aphid lineages at relatively high frequencies.

### Culturing and parallel sequencing approaches produce highly complete *Regiella* genomes

Based on the MLST screen, we selected phylogenetically diverse *Regiella* strains for genome sequencing that either belonged to a well-supported *Regiella* clade or had existing phenotypic data [26-28]. We used a combination of approaches, including a novel culturing technique with long- and short-read sequencing to produce high-quality genomes for 20 *Regiella* strains (Table 1). 5 additional strains were sequenced but removed from the study due to insufficient genome coverage (Figure S1). This approach produced highly complete chromosomal assemblies and a composite reference database of known phage and plasmid sequences to identify extrachromosomal elements. Most assemblies exhibited high CheckM completeness scores (>92.2%). Two strains of *Regiella* from *Drepanosiphum* aphids (strains 2027 and 2033) had substantially smaller genomes (1.36 Mbp) and lower GC content (39.47%). Among the assemblies, we identified 18 plasmids, clustered into 4 types, and 20 complete bacteriophages, most of which were shared among multiple *Regiella* strains (see below).

**Table 1.**
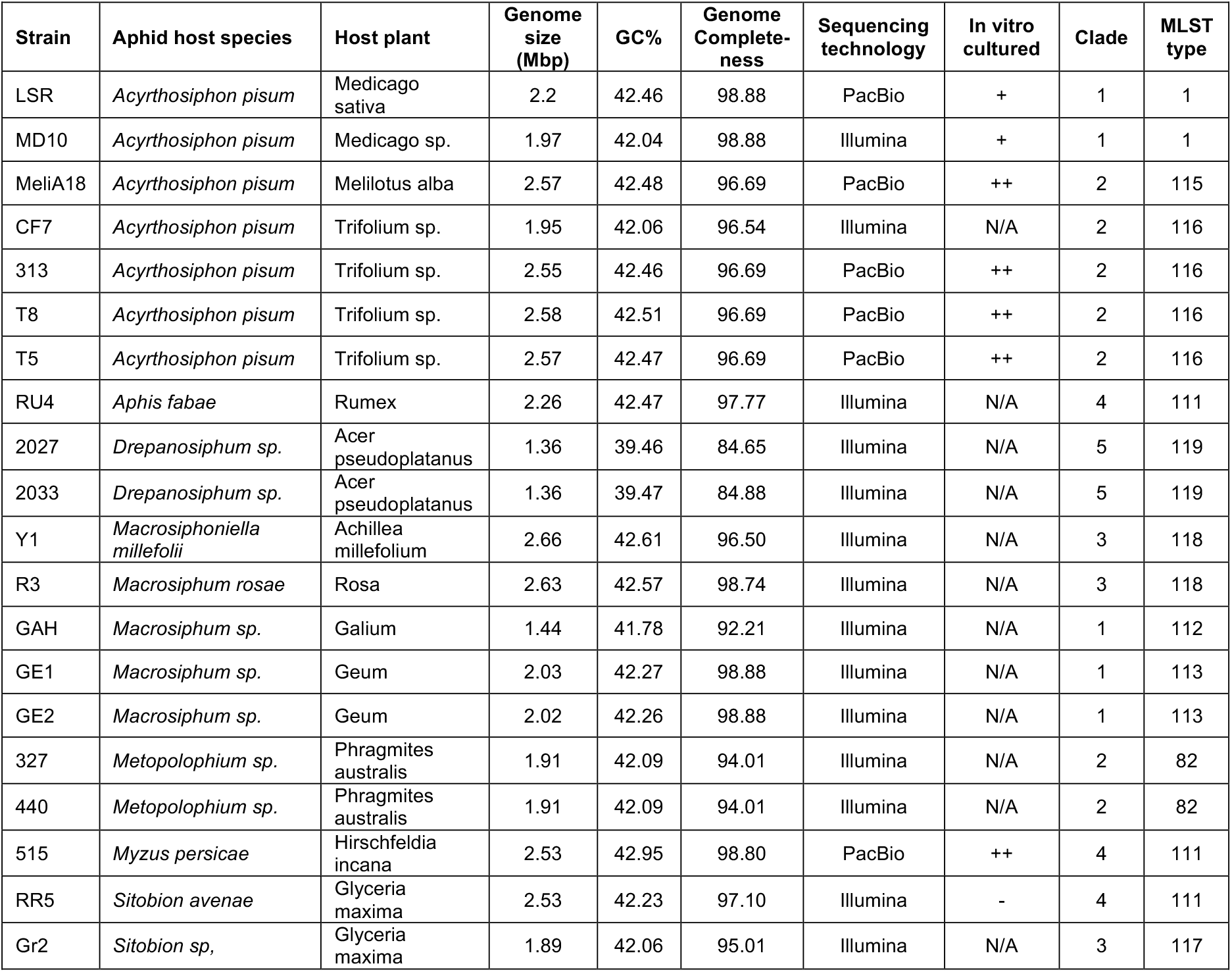
Genome statistics and metadata for *Regiella* strains analyzed in this study. For each sequenced *Regiella* genome, we indicate strain name, aphid host species, host plant from which the aphid was collected, basic genome assembly metrics including genome size, GC%, and completeness, the sequencing technology employed, *in vitro* culturing details (++ indicates adequate bacterial growth, + indicates minimal growth, - indicates no growth, N/A = not attempted), Phylogenetic clade (Matching Figure 1) and MLST type (Table S1).

### *Regiella* has diversified into multiple genetically distinct clades and has moved horizontally across aphid species

We identified five well-supported clades using a phylogeny based on 147 single-copy orthologous gene sequences from the genome assemblies (Figure 1A). Consistent with previous studies [25, 29], *Regiella* strains from different pea aphid biotypes fell into two distinct clades, and strains from these clades were also detected in other aphid species. *Regiella* from Clade 1 included strains from the *Medicago sativa* and other pea aphid biotypes (Figure S1; Table S1) grouped with strains from *Macrosiphum* species. *Regiella* stains from Clade 2, which are typically found in *Trifolium* and *Melilotus* pea aphid-biotypes [25], were also identified in *Metopolophium* aphids. These relationships, together with high amino acid sequence similarity between *Regiella* from pea aphids and non-pea aphid hosts, indicate horizontal transfer of *Regiella* among distantly related aphid species. The remaining clades, including a highly divergent lineage associated with *Drepanosiphum* aphids (which may have evolved into a co-obligate symbiont) are described in the Supplementary Results.

**Figure 1.**
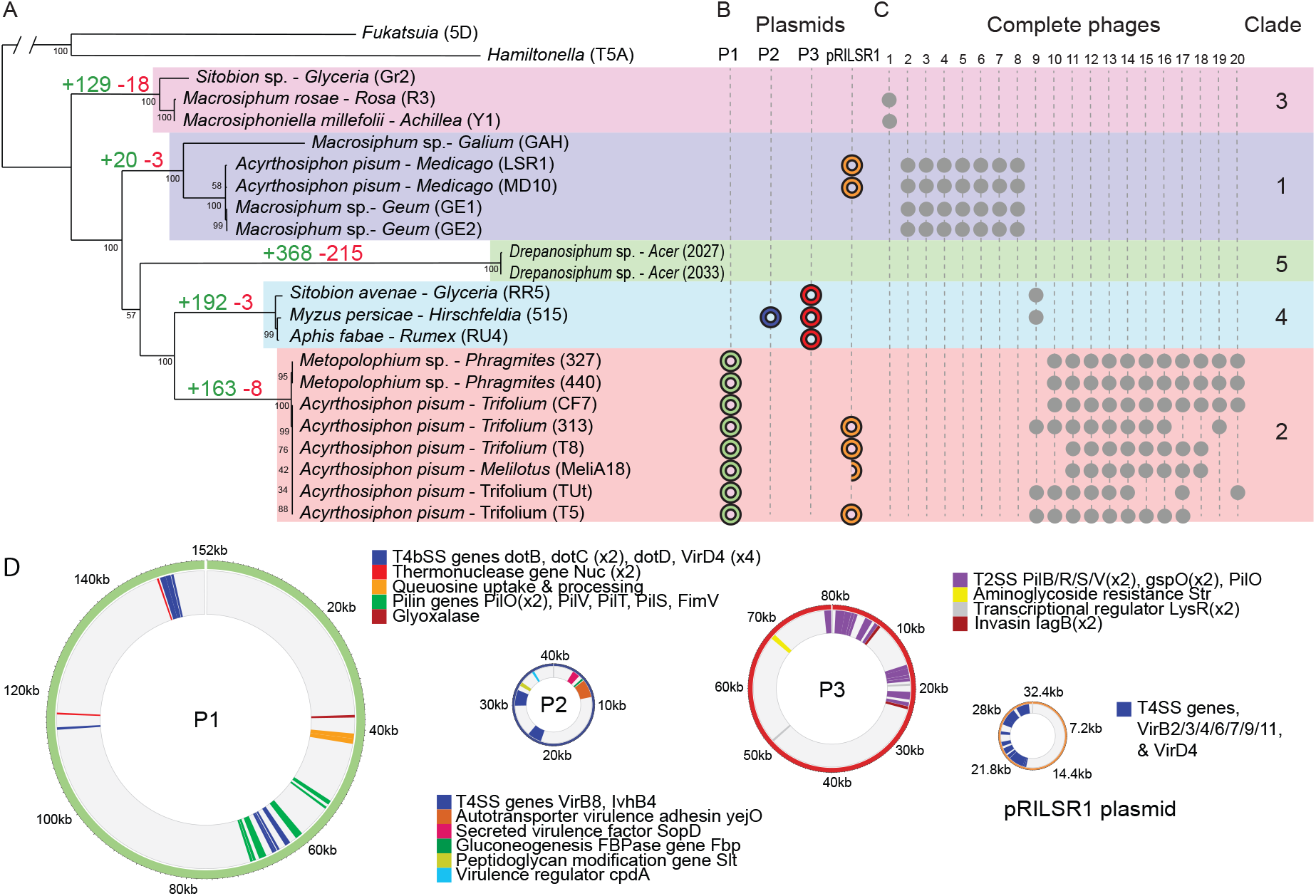
*Regiella* forms distinct phylogenetic clades with characteristic compliments of mobile genetic elements. **A**. Maximum-likelihood phylogenetic tree of all *Regiella* genome sequences based on amino acid sequences of 147 universal single copy orthologous genes. The tree is rooted to *Fukatsuia symbiotica* strain 5D (GCA_003122425.1) and *Hamiltonella defensa* strain T5A (GCA_000021705.1). Tip names represent the aphid species, the host plant genus, and the aphid clone unique identifier that each *Regiella* strain was collected from. Strains of *Regiella* are clustered into 5 distinct clades, each labelled with the number of genes uniquely present (green) or absent (red) in all strains of a particular clade. Each clade is shaded with a different color and clade numbers are indicated on the right side of the figure. **B**. Presence of plasmids in each of the *Regiella* genomes is indicated, with plasmid identity indicated along the top of the figure. **C**. Complete bacteriophages identified in *Regiella* genomes are shown. Phages were grouped according to shared coding sequences. **D**. Shows four annotated plasmids in the *Regiella* genomes, with gene categories based on annotation shown in colors as indicated in the keys.

### *Regiella* genome evolution is driven by gene gains across clades

A pangenome analysis revealed a conserved core genome shared across all sequenced strains, as well as distinct accessory gene sets that have been differentially gained or lost among clades (Figure 1A; Figure S3; Table S2). Core genome content was largely consistent across the *Regiella* phylogeny, except in strains isolated from *Drepanosiphum* aphids. We found multiple genes that were uniquely present in each clade (*i*.*e*., found only within that clade), but very few clade-specific losses (Figure 1A; Figure S4), indicating that gene gain rather than loss has had a greater impact on *Regiella’s* genome diversification. We annotated clade-specific genes using COG, KEGG, GO, and Pfam databases (Table S3; Figure S4; Supplementary Results). We found that *Regiella* clades and strains have characteristic complements of bacteriophages (Figure 1B), plasmids (Figure 1C), and transposable elements (Figure S5) that contribute gene gains across clades.

### Plasmids confer key functional gains across the *Regiella* phylogeny

Our culture-based approach enabled identification of four shared plasmids among strains based on sequence similarity (Figure 1D), most of which were missing from previous *Regiella* genome assemblies. The largest plasmid P1 (∼145kb), which contains multiple pilin genes, was exclusively found in *Regiella* clade 2 from *Trifolium* and *Melilotus* biotype pea aphids as well as *Metopolophium* aphids. This plasmid was present in the genome assembly for strain TUt [28] but was reported as part of a contig. Similarly, a second plasmid, P2 (∼40kb), was unique to the previously sequenced strain 515 (clade 4) from *Myzus* aphids and encodes Type IV secretion system genes as well as a secreted virulence factor and the virulence regulator cpdA. A third ∼80kb plasmid is present in all three clade 4 strains and encodes genes for a Type II secretion system as well as two copies of the gene IagB, which is involved in eukaryotic cell invasion and virulence. Full plasmid annotations are available in the Supplementary Material.

### The pRILSR plasmid is shared by distantly related *Regiella* clades

We found that the smallest plasmid (∼32kb, called pRILSR1), which was first reported in the *Regiella* strain LSR genome [27], is exclusively associated with *Regiella* strains from pea aphids (Figure 1A). pRILSR1 is present in *Regiella* strains from clades 1 and 2 despite their phylogenetic distance, suggesting horizontal transfer of the plasmid between *Regiella* lineages. This plasmid encodes a complete Type IV secretion system (T4SS) including VirD4 and VirB2 through VirB11. We found that pRILSR1 is absent from some pea aphid strains (CF7 and Tut) as well non-pea aphid-associated *Regiella* strains in Clades 1 and 2, including *Macrosiphum* and *Metopolophium* aphids. In addition, we found one strain (meliA18) in which a large portion of the pRILSR1 plasmid has been lost (see below).

### The pRILSR1 plasmid is implicated in fungal protection in laboratory infections

We found that even closely related *Regiella* strains differ in the fungal protection conferred to hosts (Figure 2). We established 11 *Regiella* strains derived from 3 clades into a common aphid genetic background (LSR1-01; [30]), and infected each line with the fungal pathogen *Pandora neoaphidis*. Four of the eleven strains conferred statistically significant protection against fungal infection, as indicated by a reduction in the percent of exposed aphids that produced visible sporulating cadavers compared with symbiont-free aphids. Importantly, all of the protective strains of *Regiella* harbored pRILSR1, whereas strains lacking the plasmid (as determined by genome sequencing or diagnostic PCR screening) conferred no protection (Figure 2).

**Figure 2.**
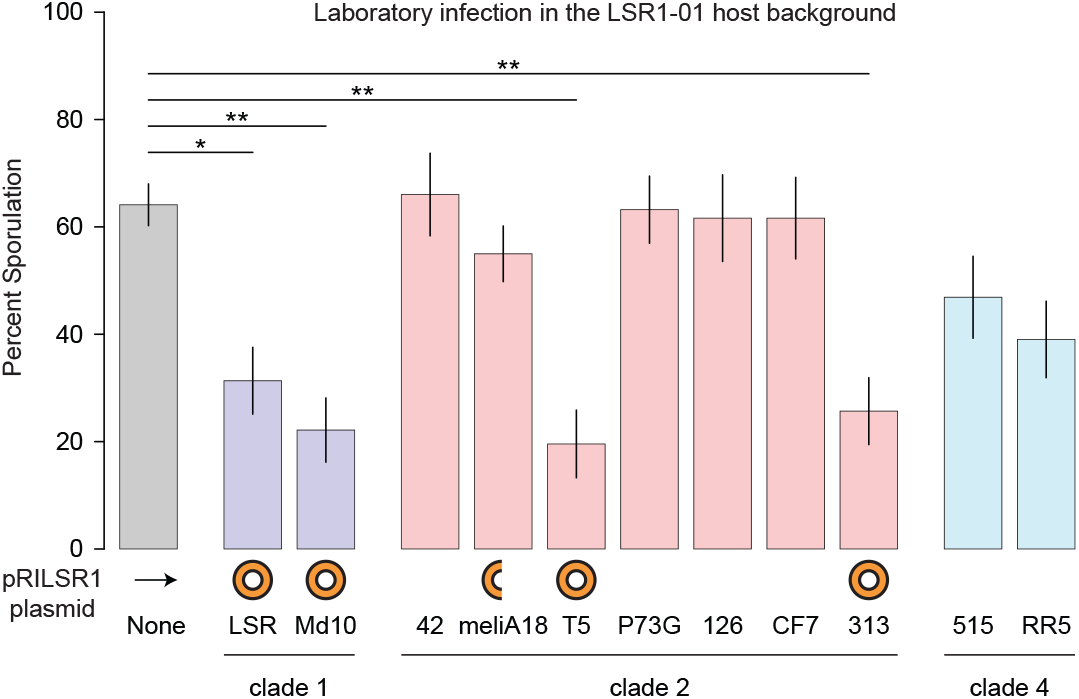
Laboratory *Pandora* infection of *Regiella* strains in a common host genetic background implicate pRILSR1 in fungal protection. The y-axis shows the percent of aphids that developed a characteristic fungal cadaver, indicating infection with *Pandora*. Symbiont strains and clades are shown along the x-axis. Control aphids without *Regiella* are shown in grey to the left of the figure. Error bars show standard error, and statistical significance is shown along the top of the figure (* < 0.05; ** < 0.01).

A genome comparison of protective and non-protective strains within *Regiella* Clade 2 demonstrates that across protective and non-protective strains, chromosomal genomes are nearly identical (T5 vs MeliA18 and 313 vs CF7 excluding the pRILSR1 plasmid = 99.99% and 99.97% AAI similarity, respectively). This indicates that the crucial difference leading to the protection is the presence of the pRILSR1 plasmid. Lastly, strain meliA18, which carried a partial copy of the pRILSR1, was not protective, demonstrating that a complete pRILSR1 plasmid is essential for *Regiella*’s fungal-protection phenotype.

### pRILSR1 is associated with fungal resistance in field-collected aphids

From a two-year field study, we found that the pRILSR1 plasmid, rather than *Regiella* presence alone, underlies fungal protection in natural populations. We collected nearly 1000 aphids from multiple host plants and recorded their survival outcomes (e.g. if they produced a *Pandora* cadaver) after 5 days in the lab. Overall, 16.4% of field aphids developed a *Pandora* infection. We screened each aphid for facultative symbionts and found, surprisingly, that at the species level *Regiella* had no significant effect on fungal infection (χ^2^ = 1.2, 924 df, p = 0.28). However, when we screened each aphid for the pRILSR1 plasmid using PCR we found that only 20.4% of *Regiella* strains carried the plasmid, and importantly, that aphids harboring pRILSR1-positive *Regiella* were significantly protected from fungal infection compared with those carrying pRILSR1-negative *Regiella* (Figure 3A; χ^2^ = 5.71, 1DF, p = 0.017). The plasmid had no effect on wasp infection (χ^2^= 0.62, 1DF, p = 0.43) or unidentified causes of mortality (χ^2^= 0.022, 1DF, p = 0.88).

**Figure 3.**
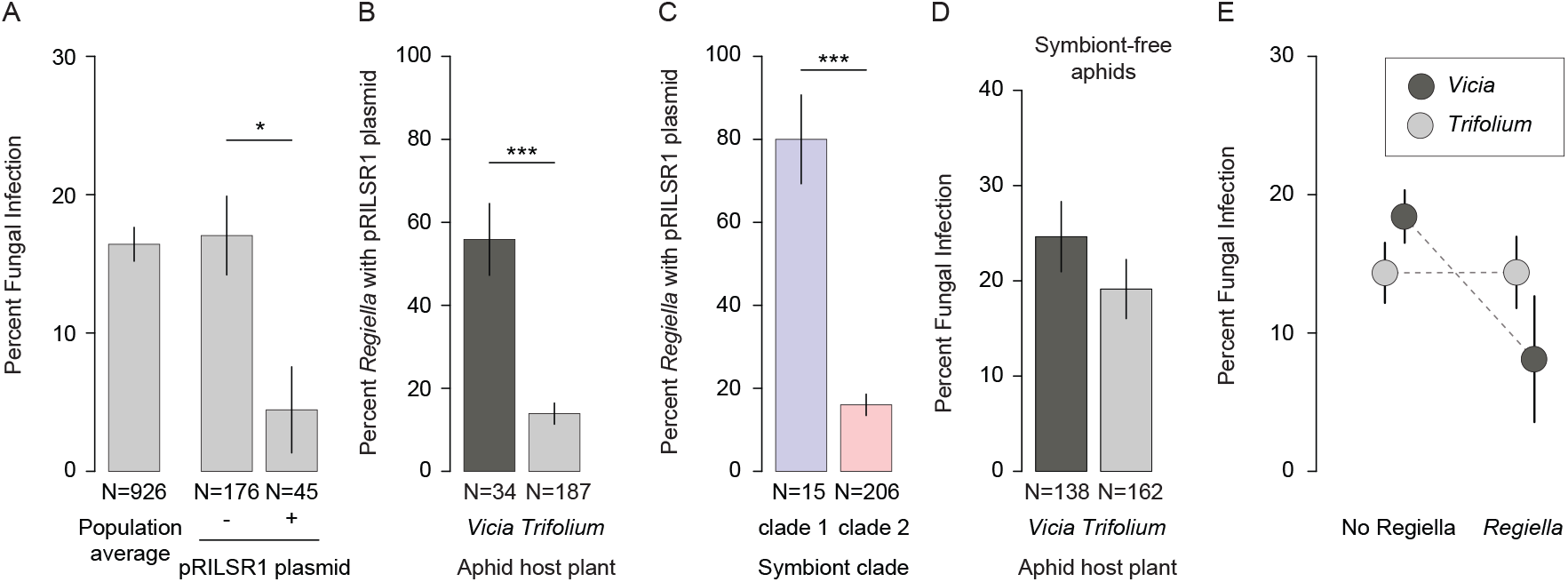
Field studies link the pRILSR1 plasmid to variation in *Pandora* protection. **A**. The y-axis shows the percent of aphids that developed a fungal cadaver when brought into the lab. Samples are grouped into the entire population (population average), aphids with *Regiella* that do not have the pRILSR1 plasmid (-), and aphids with *Regiella* that harbor the pRILSR1 plasmid (+); sample sizes are shown along the bottom of the figure. **B**. shows the percent of *Regiella* from aphids collected from *Vicia* and *Trifolium* that harbored the pRILSR1 plasmid. **C**. shows the same information for *Regiella* from clades 1 vs 2. **D**. Shows the percent of aphids that developed a fungal infection, showing only symbiont-free (no facultative symbionts of any species) from *Vicia* vs. *Trifolium*. **E**. Shows the strength of symbiont-mediated protection of each symbiont in aphids collected from *Vicia* vs. *Trifolium* host plants. The y-axis was calculated as (1 – (sporulation with symbiont / sporulation without symbiont)) *100, with complete protection against fungus conferred by a symbiont at the top of the figure and no protection at the bottom. In each panel, error bars show standard error, and statistical significance is shown along the top of the figure (* < 0.05; ** < 0.01; *** < 0.001).

### Variation in pRILSR1 frequency drives phenotypic divergence across populations

The pea aphids from our field study came from multiple host plants, including species of *Trifolium* (n = 471) and *Vicia* (n = 455). Overall, the frequency of pRILSR1 differed significantly among these pea aphid populations. It was more common in *Regiella* harbored by aphids feeding on *Vicia* than on *Trifolium* (Figure 3B; χ^2^ =16.0, 1 DF, p < 0.0001) and was also more common in clade 1 than clade 2 *Regiella* (Figure 3C; χ^2^ = 27.1, 1 DF, p < 0.0001). We found that these differences are associated with divergence in symbiont-conferred phenotypes across biotypes. Although *Pandora* prevalence did not differ significantly among symbiont-free aphids on different host plants (Figure 3D; χ^2^ = 1.32, 1 DF, p = 0.25), the impact of *Regiella* on fungal infection varied between biotypes (Figure 3E; host plant x *Regiella* interaction; χ^2^ = 3.4, 920 DF, p = 0.065), although this was only marginally significant. Together, these results show that *Vicia*-adapted pea aphids more frequently carry pRILSR1-bearing protective *Regiella*, whereas *Trifolium*-adapted aphids more often harbor non-protective *Regiella* lacking the plasmid.

### A predicted T4SS effector is highly expressed from the protective pRILSR1 plasmid

Together, these results suggest that the T4SS-encoding pRILSR1 plasmid mediates *Regiella*’s fungal protection phenotype. This is supported by the absence of virD—a critical T4SS component [31, 32]—in the partial plasmid carried by the non-protective meliA18 strain. To identify candidate T4SS effectors, we combined BastionX, a secreted-substrate predictor for Gram-negative bacteria [33], with gene-expression data from RNAseq. BastionX predicted numerous putative T4SS substrates on both the pRILSR1 plasmid and the LSR chromosome. RNAseq data from *Pandora*-infected and uninfected aphids containing the protective LSR *Regiella* stain revealed few genes were differentially expressed in response to fungal infection (Figure 4A). However, several candidate T4SS effectors were highly expressed (Figure 4B) in the *Regiella* transcriptome, and notably, the single most highly expressed gene in *Regiella* is a pRILSR1-encoded predicted T4SS effector (BastionX score 0.859; FPKM = 66,386), with an expression level nearly 100X higher than the average gene in the *Regiella* transcriptome. InterProScan identified a single conserved domain annotated as a “Secreted effector protein SifA,” and a blastp search returned hits to hypothetical proteins from diverse insect symbionts.

**Figure 4.**
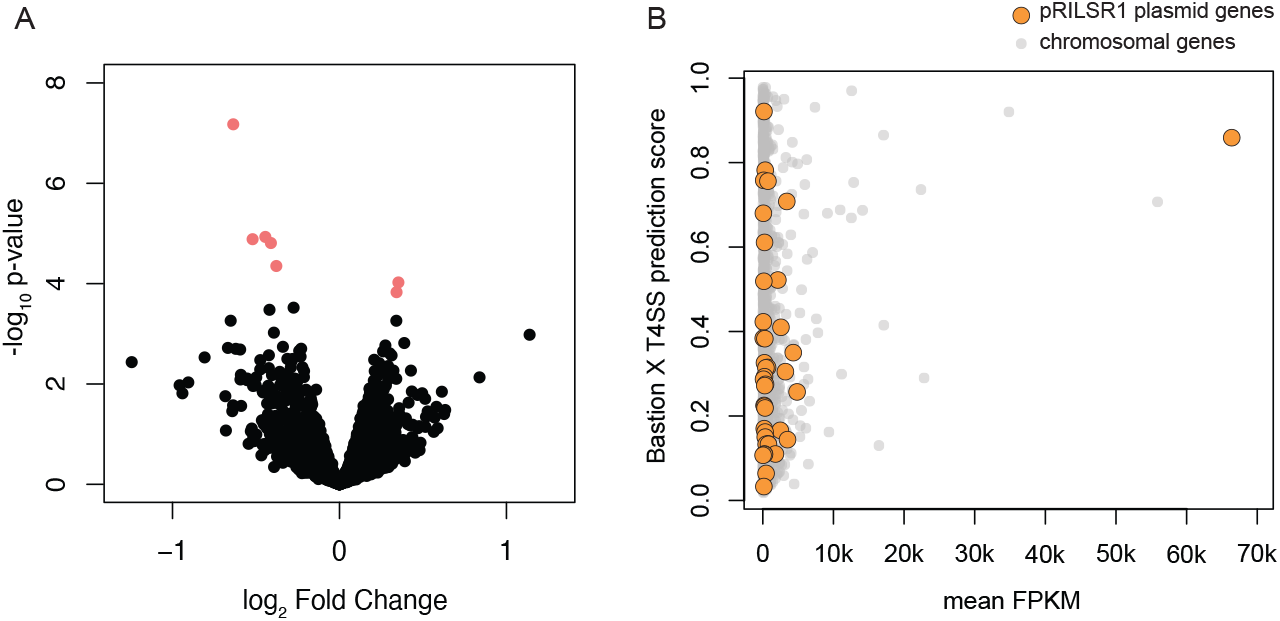
Bacterial gene expression analyses. **A**. A volcano plot comparing *Regiella* gene expression (both chromosomal and plasmid genes). Each point represents an individual bacterial gene, the x-axis shows the log 2-fold change comparing *Pandora-*infected vs. control libraries, and the y-axis shows statistical significance with genes with an FDR < 0.05 shown in red. **B**. Shows the results of a computational prediction of T4SS effectors using Bastion X, with prediction scores shown on the y-axis. The x-axis shows the strength of expression of each gene represented by FPKM. Orange points indicate pRILSR1 plasmid genes, and grey points indicate chromosomal genes.

## Discussion

Facultative symbionts can profoundly alter host biology, yet even closely related strains often confer strikingly different phenotypes. Because microbe-conferred benefits shape symbiont persistence and host adaptation, uncovering the mechanisms generating this variation is critical. Here, we show that the horizontal acquisition of a small plasmid is central to the evolution of symbiont-mediated fungal protection in aphids. The presence of this plasmid in distantly related *Regiella* lineages associated with pea aphids suggests that plasmid turnover is tightly linked to host ecology, potentially facilitated by natural co-infections of symbiont stains from different clades [29]. Our findings parallel those in *Hamiltonella defensa*, a sister-species to *Regiella*, where repeated exchange of toxin-encoding APSE phages generates strain-level differences in parasitoid resistance [34, 35]. In both systems, MGEs drive variation in defensive phenotypes, suggesting this process is a general feature that enables rapid host adaptation to variable pressures from natural enemies [36].

The pRILSR1 plasmid encodes a complete virB-like T4SS [27]. Given the established role of T4SSs in delivering effector proteins into eukaryotic cells [31, 32, 37], it is plausible that *Regiella* employs this system to secrete factors harmful to *Pandora*, either into the host hemolymph or directly into fungal cells. Effector prediction analyses identified several candidate T4SS substrates on pRILSR1, including a short protein bearing a SifA-like domain. In *Salmonella*, SifA interacts with host cellular components to promote intracellular survival and replication, suggesting an intriguing parallel, but functional validation of this candidate effector is needed. More broadly, the involvement of a secretion system in fungal protection reflects a common pattern in symbiont-mediated protection, in which mechanisms originally evolved for interacting with eukaryotic hosts are reappropriated to defend the host against eukaryotic natural enemies [38].

Variation in the benefits conferred by heritable symbionts can drive substantial phenotypic differences among host populations [11-14]. *Regiella* shows a strong and repeated association with *Trifolium*-feeding pea aphids worldwide [22-24]. Surprisingly, our field surveys show that *Regiella* does not protect *Trifolium-*adapted aphids from *Pandora*, indicating that this pattern cannot be explained by variation in fungal defense. Instead, other symbiont-encoded traits are likely contributing to the high frequency of *Regiella* on clover. Early work on the TUt strain reported improved aphid performance on *Trifolium* [22,29]. However, later studies of European *Regiella* strains found limited effect on host-plant adaptation [37,38], pointing to substantial strain-level heterogeneity. Our genomic analyses reveal that TUt harbors a distinct complement of genes lacking known homologues. Because this strain was sequenced in a previous study, we also cannot exclude the possibility that some genetic elements, such as plasmids, remain uncharacterized. Such elements could encode ecologically relevant functions, including detoxification of plant secondary compounds, enabling rapid, modular shifts in symbiont phenotypes.

Clade 2 *Regiella* also attain higher within-host densities and are better competitors, which may influence their prevenance in natural aphid populations [39]. Future genomic and functional studies should target additional *Regiella*-conferred benefits and determine whether these traits are encoded on the chromosome or on mobile genetic elements.

Recent advances in culturing and long-read sequencing are transforming our ability to resolve symbiont genomes, particularly mobile genetic elements that are often missed from earlier assemblies [34, 40].

Using these approaches, we identified MGEs missing from earlier *Regiella* assemblies, reinforcing the link between genomic plasticity and strain-specific host effects. Beyond facultative variation, our data point to a deeper evolutionary transition. Two *Regiella* strains from *Drepanosiphum* aphids form a highly divergent clade with reduced genome size, low GC content, and an accumulation of ancient transposable elements—hallmarks of genome erosion associated with co-obligate symbiosis. This pattern suggests that *Regiella* may in some lineages compensate for metabolic losses in *Buchnera*, paralleling transitions observed in other aphid systems [41]. If confirmed, this would represent a previously unrecognized case of *Regiella* transitioning toward obligate symbiosis, illustrating the dynamic evolutionary trajectories of facultative symbionts.

Strain-level variation often explains more phenotypic variation than bacterial species identity in microbial symbioses, underscoring the limitations of species-level classifications in microbial ecology [42]. This principle is likely even more pronounced in complex microbiomes, such as those of the gut, where bacterial genomes are highly dynamic due to frequent gene gain and loss mediated by mobile genetic elements. Pangenomes, rather than core genomes, often dictate functionally significant microbial traits, particularly those that influence host-microbe interactions. Our study shows how mobile genetic elements drive such variation even in the comparatively stable context of insect symbiosis. In many insect groups, long-term diversification has been shaped by symbiotic associations, with facultative symbionts contributing to ecological adaptation. Together, our results highlight mobile genetic elements as a key resource driving rapid, context-dependent evolution of host-benefiting symbiont traits and phenotypic divergence among insect populations.

## Methods

### Sample collection for analysis of *Regiella* genetic diversity

We collected aphids in the UK from 2011 to 2020 from multiple host plants and we preserved samples in ethanol. We extracted genomic DNA from individual aphid samples using DNeasy Blood and Tissue kits (QIAGEN, Venlo, Netherlands) under recommended conditions. To identify aphid species, we amplified an approximately 700 bp DNA fragment of the cytochrome c oxidase I (COI) ‘barcoding’ gene from the DNA using the Lep_F and Lep_R primers [43], and Sanger sequenced the PCR products in the forward direction. We aligned DNA sequences with MUSCLE [44] and identified aphid species by comparing COI sequences to the BOLD database (http://www.boldsystems.org/) and GenBank using BLASTn [45].

### Screening and *Regiella* genotyping

We screened aphid genomic DNA for *Regiella* using diagnostic PCR of the 16S ribosomal RNA gene [46]. For samples containing *Regiella*, we used a multilocus sequence typing (MLST) scheme to characterize *Regiella* lineages based on five housekeeping genes: *accD, gyrB, murE, recJ*, and *rpoS* [46]. MLST amplicons were Sanger-sequenced in the forward direction. We used Tracy [47] for base calling on the chromatograms to confirm nucleotide substitutions, and we aligned concatenated MSLT genes using MUSCLE with default parameters. Phylogenetic analyses (see below) were conducted on the concatenated MLST alignment.

### *Regiella* culturing, DNA extraction, and genome sequencing

We established *in vitro* cultures of *Regiella* strains LSR, MD10, MeliA18, 313, T8, T5, and 515 in insect cell lines [9]. These strains all originated from pea aphids (*A. pisum*) with the exception of strain 515, which was isolated from *Myzus persicae* and then transferred into pea aphids for lab colony maintenance and initiating the *in vitro* culture. We attempted to establish strain RR5 in culture, but this produced no visible growth. We extracted DNA from bacterial pellets as described in previous work [48]. Purified genomes of *Regiella* were sequenced on a PacBio Sequel II platform at an average yield of 150,000 reads of 10kbp length per sample. Strain LSR was sequenced using an ultralow input sample prep method, giving a yield of 1.4 million reads of 10kbp length. We also performed sequencing on 12 additional strains of *Regiella* (Table 1) using gDNA extracted with the EZNA Insect DNA kit (Omega Bio-Tek, Norcross, Ga, USA) with an Illumina NovaSeq S4 following the NEBNext Ultra II FS kit library prep protocol to generate paired-end reads of 2x150bp length at an average of 200 million reads per sample. These samples were sequenced at the Centre for Genomic Research at the University of Liverpool. A final sample (strain CF7) was sequenced on DNA extracted using the Qiagen DNEasy kit at Novogene, Inc., on an Illumina NovaSeq PE150 platform to 15 Gbp.

### Genome assembly

We checked reads for quality and adapter content using FastQC and MultiQC [49] and used FastP [50] with default parameters for trimming. We assembled the PacBio long reads using MetaFLYE v2.9 [51] with parameters --pacbio_hifi and 5 polishing iterations. We assembled the Illumina reads in two steps. First we used SPAdes v3.15.4 [52] in --assembly-only mode with default parameters. We then mapped the reads to the assembly using BWA-mem [53] and assigned taxonomic identities to the contigs using megablast [54] and DIAMOND [55] searches against the NCBI’s non-redundant Refseq nucleotide and protein databases respectively. We used BlobTools v1.1.1 [56] to examine and extract the contigs matching *Regiella* showing distinct coverage and GC content. Then, we extracted *Regiella* reads with SAMtools [57] and reassembled them separately to give more contiguous and complete genomes with SPAdes v3.15.4 [52] in --careful mode using error correction and kmer sizes of 33, 55, 77, 99, and 127.

We manually inspected all final assembly graphs in Bandage [58] for uniform coverage and contiguity, and we performed BlastN [45] searches against the NCBI nonredundant nucleotide database to validate the taxonomic identities of the contigs. We assessed the completeness and contamination levels of the assemblies with CheckM [59] using f_Enterobacteriaceae as the marker lineage that has 546 genomic marker genes. We checked the genomic markers and functional genes that were automatically assigned as missing using HMM [60] and blast [45] searches of the target genes on the raw contigs from the whole-aphid initial assembly. To validate the presence of plasmids in strains sequenced with short reads, we used the contiguous circularized plasmid sequences from the PacBio sequenced strains, as well as 25 previously identified plasmids from *Regiella, Hamiltonella*, and *Fukatsuia* in addition to the nucleotide sequences of all known plasmid incompatibility (Inc) groups to create a BLASTn nucleotide database. This database was used to perform BLASTn searches of all the contigs generated by short read sequencing to query the Illumina sequenced strains for plasmids. Since many of these plasmids share similar genes for their backbones and plasmid maintenance, a strict threshold of >80% coverage was used to robustly identify and indicate the presence of plasmids in the short read sequenced samples.

### Phylogenetic analyses

We used PhyML [61] with default parameters and 500 bootstraps to reconstruct the *Regiella* MLST phylogeny, and ITOL [62] to visualize the phylogenetic tree. To construct the genome-based phylogeny of all the assembled *Regiella* strains and the previously sequenced Tut strain (GCA_013373955.1), we ran the GToTree v1.8.2 [63] pipeline using HMMER3 v3.3.2 [60], Muscle 5.1.linux64 [44], TrimAL v1.4.rev15 [64], Prodigal v2.6.3 [65], ModelFinder [66], UFBoot2 [67], GNU Parallel v20230922 [68], and IQ-Tree v2.2.5 [69] in default settings with 1000 bootstraps in RAxML v8 [70] on a concatenated alignment of the 172 orthologous genes universally present in Gammaproteobacteria. We visualized the phylogeny and rooted it to *Fukatsuia symbiotica* strain 5D (GCA_003122425.1) and *Hamiltonella defensa* strain T5A (GCA_000021705.1) in ITOL [62].

### Pan-genome analysis

We used Prokka v1.14.6 with default settings in --compliant mode to annotate all the assembled genomes [71]. We performed a new annotation the genome of strain Tut to ensure uniformity of the annotations. We used Roary v3.13 [72] with -e -i 90 -s settings to identify core genes common at 90% amino acid identity cutoff in all genomes and accessory genes unique to *Regiella* strains. We visualized the pangenome matrix using Phandango [73]. We annotated the genes uniquely present and absent in each *Regiella* clade with COG, KEGG, GO terms, and Pfam domains using eggnog v6.0 [74]. We visualized the annotated functional groups with GGplot2 in R v4.3.2. through Rstudio (33206f75, 2023-12-17).

### Comparative analysis of gene function

We further analyzed the genomes to explore the functional categories that had been gained in each *Regiella* clade. We used Phaster [75] on default parameters to identify and extract bacteriophage sequences from the assemblies. We filtered out incomplete sequences, and analyzed the complete bacteriophage sequences using PhageScope [76] to determine their taxonomy and generate genome statistics. We identified transposable elements and analyzed them using Earl Grey [77]. We identified the presence of genes coding for the synthesis of amino acids, vitamins, carbon, lipid, and nucleotide metabolism and transporters using KofamKoala [78] and analyzed them in KEGGMapper [79]. Last, we visualized the protein coding genes that are on plasmid and have a function other than the transmission or replication of plasmids using Circos plots [80]. To validate the absence of genes uniquely missing in clades, we manually checked for these genes in the raw assemblies to ensure reported absences were not due to differences in genome coverage or sequencing technologies used.

### Experimental fungal infections

We transfected 11 *Regiella* strains from 3 clades into a common pea aphid host genetic background (LSR1-01; [30]), and waited at least 4 generations for the symbiosis to stabilize as in previous studies [81]. We infected aphids with *Pandora neoaphidis* using previously described methods [82]. Briefly, we exposed aphids to sporulating fungal cadavers in a PVC tube infection chamber, with sporulating cadavers placed above the chamber so spores fall onto the experimental aphids. Sporulating cadavers were rotated among the treatment aphids so that all aphids within an experiment are exposed to each set of sporulating cadavers for an equal period of time as in [83]. We then transferred aphids to a fava bean plant (5-10 aphids per plant) with a sealed cup cage over the plant, which produces humid conditions necessary for *Pandora* to infect aphids. We kept aphids and plants at 20°C and 16L:8D for 48 hours, after which we transferred aphids to a new plant without parafilm. We recorded whether each aphid survived, died of unknown causes, or produced a sporulating cadaver every 24hrs. For the experimental infection assay of *Regiella* strains, we used a *Pandora* isolate collected in Knoxville, TN in 2019 established from a pea aphid collected from *Vicia*. For the RNAseq assay, we used a *Pandora* isolate collected in Chapel Hill, NC in 2025 from *Macrosiphum euphorbiae* feeding on tomato. Both strains were confirmed to be *Pandora neoaphidis* using PCR amplification and Sanger sequencing of the ITS gene as in previous work [84].

### Field study collections and sample storage

We collected adult pea aphids in consecutive springs (2021-2022) in Knox County, TN, USA feeding on species from two genera of host plants in the family Fabaceae (*Trifolium campestre, T. incarnatum, T. pratense, T. repens, Vicia grandiflora, V. sativa, V. villosa*,). Aphids were collected from seven sites with large swaths of host plants: Tommy Schumpert Park (160 acres; N 36.046, W -83.958), Cherokee Farms Park (147 acres; GPS: N 35.946, W -83.952), Baker Creek Park (23 acres; N 35.942, W -83.890), I.C. King Park (10 acres; N 35.903, W -83.940), Tyson McGee Park (13 acres; N 35.951, W -83.942), Knoxville Botanical Garden (47 acres; N 35.982, W -83.883), and Henry Ave (5 acres; N 35.941, W -83.912). We visited these sites weekly from April 6 through June 23, 2021 (87 days), and in 2022 from March 30 through June 14 (78 days).

To collect aphids, we placed a plastic tray under each host plant and hit the stem with a stick to dislodge feeding aphids. We then placed one adult pea aphid from the tray in a Petri dish (100mm) with a terminal end cutting of the host plant from which it was collected. The location, plant species, and date were recorded. Back in the lab, we poured a small amount (∼3mL) of tap water agar into each Petri dish to secure the plant cutting and prevent desiccation. Each dish was then sealed with parafilm. We kept each dish on a lab bench at room temperature for five days, and then scored each aphid phenotype as remaining healthy, dead with a visible fungal infection (a fungal ‘cadaver’), dead with a visible wasp parasitism resulting in aphid mummification, or dead from an unknown cause. We then transferred each aphid to a microcentrifuge tube and stored at -20ºC. If present, up to five clonal offspring from each adult were also stored in a separate tube.

### Field sample DNA extraction and screening

We extracted DNA from 926 adult aphids using ‘Bender buffer’ as in [85]. In short, we crushed frozen aphids with sterile pestles in microcentrifuge tubes and heat-treated (2 hours at 65°C) each sample with the lysis buffer and Proteinase K (Fisher Bioreagents). We subsequently treated samples with 8M potassium acetate, vortexed each tube, and kept extractions on ice overnight followed by three ethanol washes and centrifugations. Following precipitation, we eluted the DNA pellet in nuclease-free water and diluted each sample to 20ng/μL.

We carried out separate diagnostic PCR screens for seven bacterial symbionts: *Regiella insecticola, Hamiltonella defensa, Serratia symbiotica, Spiroplasma* sp., *Rickettsia* sp., *Ricketsiella viridis*, and *Fukatsuia symbiotica*. We used the following PCR program: 94°C 5min, 11 cycles of 94°C 30s, 56°C (declining 1°C each cycle) 50s, 72°C 90s, 25 cycles of 94°C 30s, 45°C 50s, 72°C 90s and a final extension of 72°C 5min as in [25]. For each PCR reaction, extracted DNA from known symbiont-containing aphids was used as positive controls, while sterile water was used as the negative control. We extracted DNA from clonal offspring of a subset of samples and screened for symbionts to assess agreement between adult and clone symbiont communities, and we found strong concurrence between PCR reactions from adult fungal cadavers and offspring.

### Strain typing of field *Regiella*

To characterize the genetic diversity of *Regiella* field samples, we used two methods. For the majority of samples, we amplified and sequenced the murE housekeeping gene specific to *Regiella* containing samples using a primer pair and PCR recipe as above [25, 46]. Amplicons were gel extracted and purified (Zymo Clean and Concentrator Kit under recommended conditions) and sequenced in forward and reverse directions using Sanger sequencing. A representative sample from each clade was used as a PCR positive control as well as for sequencing. Consensus sequences of experimental samples and representative samples (LSR – clade 1; 313 – clade 2) were generated by aligning forward and reverse sequences with ends trimmed by hand. A maximum likelihood phylogeny was generated as above to assign strains to each clade. For some samples we amplified a polymorphic region of the *Regiella* genome where some strains have a 32bp deletion in the amplicon as in [29].

Amplicons were visualized on a 3.5% agarose gel run at 70V on ice for 3.5 hours to reduce any gel streaking and confirm the size of amplicons and assign remaining samples to clade 1 or 2. Samples that failed both PCR screens for *Regiella* diversity (n=6) were included in results after re-screening with original symbiont check PCR and removing false positives.

### PCR screening for pRILSR1 plasmid

We designed primers that amplified a 701bp region of the pRILSR1 plasmid (133F 5’-CAACGTATCAGCGCCAATGG-3’ and 834R 5’-CGGCTCGATGCTGTTTTCTG-3’). We used NEB 2X Quick Load Master mix to run 20μL reactions that included 10μL of master mix, each primer at 10μM, and 40ng of gDNA. PCR conditions included an initial step of 95°C for 3m, 40 cycles of 95°C for 30s, 56.5°C for 30s and 72°C for 60s, followed by a final extension step of 72°C for 5m and a 12°C hold. We screened each of the *Regiella* positive samples for the presence or absence of the plasmid. We analyzed the percent of *Regiella* from each clade and each host plant with the plasmid and the effect of plasmid carriage on fungal infection using generalized linear models with a binomial error structure, comparing models using Chi squared tests, implemented in R v.4.4.3 [86].

### T4SS effector prediction and RNAseq

For T4SS effector prediction, the *Regiella* strain LSR genome annotation (from this study) was used to extract the protein sequences for each predicted gene using gffread. We uploaded these to Bastion Hub (https://bastionhub.erc.monash.edu/index.jsp) and analyzed sequences using Bastion X V2.0 on the highest sensitivity settings. To look for conserved domains in candidate genes, we used Interpro v 106.0, using InterProScan 5.75-106.0. For RNAseq, we collected aphids from the LSR1-01 background harboring strain LSR at 60hrs after fungal exposure, and stored samples at -80°C. We also included control samples at the same timepoint that were handled similarly but not exposed to fungus. We selected 60h because previous work has shown that the fungus takes between 24 and 48 hours to cross the aphid cuticle [87], but by 72hrs has converted much of the aphid’s biomass to mycelium as it prepares to produce a sporulating cadaver between 72hrs and 96hrs post infection. We extracted RNA from each sample by macerating whole aphids in groups of 3 individuals per treatment per timepoint in 300 μL Trizol (Invitrogen; Thermo Fisher Scientific, Inc.,) with 100 μL BCP (1-bromo-3-chloropropane; Life Technologies, Thermo Fisher Scientific, Inc.) and standard isopropanol precipitation. We resuspended each RNA pellet in 40 μL nuclease-free water, removed genomic DNA with the Zymo DNase I Reaction Mix according to the manufacturer instructions (Zymo Genetics, Inc.,) and cleaned and concentrated with the Zymo RNA Clean & Concentrator kit under recommended conditions. Library preparation of 4 biological replicates per treatment including ribosomal RNA removal and sequencing was carried out by Novogene Inc. We mapped reads to a combined fasta file containing pea aphid genome, *Buchnera aphidicola* strain LSR genome, and the LSR genome assembly from above using BWA v.0.7.17 [53], and counted reads aligned to the LSR genome with htseq v.2.0.5 [88].

We then assessed differential gene expression using edgeR v.3.40.2 [89]. We calculated average FPKM across the 8 libraries as read count / (gene length in kb x library size).

## Acknowledgements

This work was funded in part by a UKRI/BBSRC-NSF/BIO Lead Agency Opportunity award to L.M.H (BBSRC grant BB/W001632/1) and B.J.P. (U.S. National Science Foundation Grant IOS-2152954), and in part by NIH/NIGMS under award number R35GM154800 to B.J.P. M.K. was supported by NSF Postdoctoral Research Fellowship in Biology award DBI-2109440.

B.J.P. is a Pew Scholar in the Biomedical Sciences, funded by the Pew Charitable Trusts. Joseph Torres helped with bioinformatic analyses, Ashlyn Anderson with nucleic acid extraction, and William Brewer, Holly Nichols, Taylor Do, McKayla Martin, Georgina Aitolo, Lance Knipper, and Sudan Maharjan helped

## Supplementary Tables and Figures

**Table S1:** Metadata of collected *Regiella*-positive aphids, aphid genus, host plant, and sequence type (ST) of *Regiella* strains based on 5 MLST genes.

**Table S2:** Pangenome comparison table of all genes found in all analyzed genomes of *Regiella* strains. Columns indicate the different annotation assignments for each gene tagged with their respective common gene name, description of the gene product, COG category, GO annotation, EC identifier, KEGG KO number annotation, and Pfam domain identifier.

**Table S3:** Annotations of genes uniquely present and absent in each phylogenetic clade of *Regiella*.

**Table S4:** Annotations of genes present in the 20 different strains of bacteriophages carried by *Regiella*.

**Table S5:** Complete bacteriophages identified in *Regiella* genomes, GC content, phage size, predicted host, lifestyle, and taxonomic classification.

**Table S6:** Annotations of protein coding genes on plasmids.

**Table S7:** Field data.

**Table S8:** Statistical analysis of symbiont co-infections in field data.

**Figure S1:** Maximum-likelihood phylogeny of field collected *Regiella* strains using five housekeeping genes. Bars next to each node indicate the proportions of the aphid host species of the symbionts. Branches labelled with green indicate the representative strains used for whole genome sequencing. Additional strains from in-house cultures (MD10, CF7, T8, T5, 2027, R3, GE1, 440, 515, RR5) were also sequenced. Five strains that were sequenced (GA2, U3, Y2, 326, MeliA11) were removed from the study due to insufficient genome coverage.

**Figure S2:** Maximum-likelihood phylogeny of the Cytochrome Oxidase 1 gene copies of the *Regiella*-positive aphids sampled in this study. Bars next to each node indicate the genotype of the *Regiella* strains associated with each aphid strain. *Regiella* strains with more than 99% nucleotide identity were collapsed to single genotypes.

**Figure S3:** Phylogeny and pangenome matrix of all *Regiella* genomes. Presence/absence of genes shared highlight the genetic diversity across different strains

**Figure S4:** Counts of functional categories of annotated genes uniquely present/absent in each clade of *Regiella*. Functional categories are based on the manual curation of aggregated annotation results from multiple tools GO, COG, KEGG, and Pfam.

**Figure S5:** Matrix of transposable elements (TEs) in *Regiella* genomes. TEs are grouped into families as orthogroups (top) and dot sizes indicate the number of TEs per strain in each orthogroup.

**Figure S6:** A) Proportion of genomes covered by TEs. B) Repeat landscape plots of TEs. Y-axis shows the percent abundance of each TE family, X-axis shows the kimura distance between TEs of each family used as a proxy for relative time of activity in the genome.

**Figure S7:** Pangenome matrix of all complete bacteriophages present in the *Regiella* genomes. Phylogeny indicates the relatedness of phages from different strains of *Regiella* based on the number of phage genes shared

**Figure S8:** Symbiont infection frequencies and patterns of co-infection in the Tennessee field experiment.

**Figure S9:** Temporal patterns in the Tennessee field experiment.

## References

1. Buchner P. Endosymbiosis of animals with plant microorganisms (revised english version). New York: John Wiley & Sons, Inc.; 1965.

2. Perreau J, Moran NA. Genetic innovations in animal-microbe symbioses. Nat Rev Genet. 2022;23(1):23–39. Epub 20210813. doi: 10.1038/s41576-021-00395-z. PubMed PMID: 34389828; PubMed Central PMCID: PMCPMC8832400.

3. Tokuda G, Mikaelyan A, Fukui C, Matsuura Y, Watanabe H, Fujishima M, et al. Fiber-associated spirochetes are major agents of hemicellulose degradation in the hindgut of wood-feeding higher termites. Proc Natl Acad Sci U S A. 2018;115(51):E11996–E2004. Epub 20181130. doi: 10.1073/pnas.1810550115. PubMed PMID: 30504145; PubMed Central PMCID: PMCPMC6304966.

4. Bennett GM, Moran NA. Heritable symbiosis: The advantages and perils of an evolutionary rabbit hole. Proc Natl Acad Sci U S A. 2015;112(33):10169–76. Epub 2015/02/26. doi: 10.1073/pnas.1421388112. PubMed PMID: 25713367; PubMed Central PMCID: PMCPMC4547261.

5. Siozios S, Nadal-Jimenez P, Azagi T, Sprong H, Frost CL, Parratt SR, et al. Genome dynamics across the evolutionary transition to endosymbiosis. Curr Biol. 2024. Epub 20241112. doi: 10.1016/j.cub.2024.10.044. PubMed PMID: 39549700.

6. Jaenike J. Population genetics of beneficial heritable symbionts. Trends Ecol Evol. 2012;27(4):226–32. Epub 2011/11/23. doi: 10.1016/j.tree.2011.10.005. PubMed PMID: 22104387.

7. Gu X, Ross PA, Gill A, Yang Q, Ansermin E, Sharma S, et al. A rapidly spreading deleterious aphid endosymbiont that uses horizontal as well as vertical transmission. Proc Natl Acad Sci U S A. 2023;120(18):e2217278120. Epub 20230424. doi: 10.1073/pnas.2217278120. PubMed PMID: 37094148; PubMed Central PMCID: PMCPMC10161079.

8. Moran NA, McCutcheon JP, Nakabachi A. Genomics and evolution of heritable bacterial symbionts. Annu Rev Genet. 2008;42:165–90. Epub 2008/11/06. doi: 10.1146/annurev.genet.41.110306.130119. PubMed PMID: 18983256.

9. Brandt JW, Chevignon G, Oliver KM, Strand MR. Culture of an aphid heritable symbiont demonstrates its direct role in defence against parasitoids. Proc Biol Sci. 2017;284(1866). Epub 2017/11/03. doi: 10.1098/rspb.2017.1925. PubMed PMID: 29093227; PubMed Central PMCID: PMCPMC5698653.

10. Masson F, Calderon Copete S, Schupfer F, Garcia-Arraez G, Lemaitre B. In Vitro Culture of the Insect Endosymbiont Spiroplasma poulsonii Highlights Bacterial Genes Involved in Host-Symbiont Interaction. MBio. 2018;9(2). Epub 2018/03/22. doi: 10.1128/mBio.00024-18. PubMed PMID: 29559567; PubMed Central PMCID: PMCPMC5874924.

11. Martinez J, Tolosana I, Ok S, Smith S, Snoeck K, Day JP, et al. Symbiont strain is the main determinant of variation in Wolbachia-mediated protection against viruses across Drosophila species. Mol Ecol. 2017;26(15):4072–84. Epub 2017/05/04. doi: 10.1111/mec.14164. PubMed PMID: 28464440; PubMed Central PMCID: PMCPMC5966720.

12. McLean AHC, Hrcek J, Parker BJ, Mathe-Hubert H, Kaech H, Paine C, et al. Multiple phenotypes conferred by a single insect symbiont are independent. Proc Biol Sci. 2020;287(1929):20200562. Epub 2020/06/18. doi: 10.1098/rspb.2020.0562. PubMed PMID: 32546097; PubMed Central PMCID: PMCPMC7329050.

13. Heyworth ER, Ferrari J. A facultative endosymbiont in aphids can provide diverse ecological benefits. J Evol Biol. 2015;28(10):1753–60. Epub 2015/07/25. doi: 10.1111/jeb.12705. PubMed PMID: 26206380; PubMed Central PMCID: PMCPMC4949989.

14. Nikoh N, Hosokawa T, Moriyama M, Oshima K, Hattori M, Fukatsu T. Evolutionary origin of insect-Wolbachia nutritional mutualism. Proc Natl Acad Sci U S A. 2014;111(28):10257–62. Epub 2014/07/02. doi: 10.1073/pnas.1409284111. PubMed PMID: 24982177; PubMed Central PMCID: PMCPMC4104916.

15. Douglas AE. NUTRITIONAL INTERACTIONS IN INSECT-MICROBIAL SYMBIOSES: Aphids and Their Symbiotic Bacteria Buchnera. Annual Review of Entomology. 1998;43:17–37.

16. Oliver KM, Degnan PH, Burke GR, Moran NA. Facultative symbionts in aphids and the horizontal transfer of ecologically important traits. Annu Rev Entomol. 2010;55:247–66. Epub 2009/09/05. doi: 10.1146/annurev-ento-112408-085305. PubMed PMID: 19728837.

17. Parker BJ, Spragg CJ, Altincicek B, Gerardo NM. Symbiont-mediated protection against fungal pathogens in pea aphids: a role for pathogen specificity? Appl Environ Microbiol. 2013;79(7):2455–8. Epub 2013/01/29. doi: 10.1128/AEM.03193-12. PubMed PMID: 23354709; PubMed Central PMCID: PMCPMC3623210.

18. Scarborough CL, Ferrari J, Godfray HC. Aphid protected from pathogen by endosymbiont. Science. 2005;310(5755):1781. Epub 2005/12/17. doi: 10.1126/science.1120180. PubMed PMID: 16357252.

19. Vorburger C, Gehrer L, Rodriguez P. A strain of the bacterial symbiont Regiella insecticola protects aphids against parasitoids. Biol Lett. 2010;6(1):109–11. Epub 2009/09/25. doi: 10.1098/rsbl.2009.0642. PubMed PMID: 19776066; PubMed Central PMCID: PMCPMC2817266.

20. Higashi CHV, Nichols WL, Chevignon G, Patel V, Allison SE, Kim KL, et al. An aphid symbiont confers protection against a specialized RNA virus, another increases vulnerability to the same pathogen. Mol Ecol. 2023;32(4):936–50. Epub 20221220. doi: 10.1111/mec.16801. PubMed PMID: 36458425; PubMed Central PMCID: PMCPMC10107813.

21. Tsuchida T, Koga R, Fukatsu T. Host plant specialization governed by facultative symbiont. Science. 2004;303(5666):1989. Epub 2004/03/27. doi: 10.1126/science.1094611. PubMed PMID: 15044797.

22. Ferrari J, West JA, Via S, Godfray HC. Population genetic structure and secondary symbionts in host-associated populations of the pea aphid complex. Evolution. 2012;66(2):375–90. Epub 2012/01/27. doi: 10.1111/j.1558-5646.2011.01436.x. PubMed PMID: 22276535.

23. Russell JA, Weldon S, Smith AH, Kim KL, Hu Y, Lukasik P, et al. Uncovering symbiont-driven genetic diversity across North American pea aphids. Mol Ecol. 2013;22(7):2045–59. Epub 2013/02/06. doi: 10.1111/mec.12211. PubMed PMID: 23379399.

24. Tsuchida T, Koga R, Shibao H, Matsumoto T, Fukatsu T. Diversity and geographic distribution of secondary endosymbiotic bacteria in natural populations of the pea aphid, Acyrthosiphon pisum. Mol Ecol. 2002;11(10):2123–35. Epub 2002/09/26. doi: 10.1046/j.1365-294x.2002.01606.x. PubMed PMID: 12296954.

25. Henry LM, Peccoud J, Simon JC, Hadfield JD, Maiden MJ, Ferrari J, et al. Horizontally transmitted symbionts and host colonization of ecological niches. Curr Biol. 2013;23(17):1713–7. Epub 2013/09/03. doi: 10.1016/j.cub.2013.07.029. PubMed PMID: 23993843; PubMed Central PMCID: PMCPMC3980636.

26. Hansen AK, Vorburger C, Moran NA. Genomic basis of endosymbiont-conferred protection against an insect parasitoid. Genome Res. 2012;22(1):106–14. Epub 2011/09/29. doi: 10.1101/gr.125351.111. PubMed PMID: 21948522; PubMed Central PMCID: PMCPMC3246197.

27. Degnan PH, Leonardo TE, Cass BN, Hurwitz B, Stern D, Gibbs RA, et al. Dynamics of genome evolution in facultative symbionts of aphids. Environ Microbiol. 2010;12(8):2060–9. Epub 2010/08/01. doi: 10.1111/j.1462-2920.2009.02085.x. PubMed PMID: 21966902; PubMed Central PMCID: PMCPMC2955975.

28. Nikoh N, Tsuchida T, Koga R, Oshima K, Hattori M, Fukatsu T. Genome Analysis of “Candidatus Regiella insecticola” Strain TUt, Facultative Bacterial Symbiont of the Pea Aphid Acyrthosiphon pisum. Microbiol Resour Announc. 2020;9(40). Epub 20201001. doi: 10.1128/MRA.00598-20. PubMed PMID: 33004445; PubMed Central PMCID: PMCPMC7530917.

29. Guyomar C, Legeai F, Jousselin E, Mougel C, Lemaitre C, Simon JC. Multi-scale characterization of symbiont diversity in the pea aphid complex through metagenomic approaches. Microbiome. 2018;6(1):181. Epub 2018/10/12. doi: 10.1186/s40168-018-0562-9. PubMed PMID: 30305166; PubMed Central PMCID: PMCPMC6180509.

30. International Aphid Genomics C. Genome sequence of the pea aphid Acyrthosiphon pisum. PLoS Biol. 2010;8(2):e1000313. Epub 2010/02/27. doi: 10.1371/journal.pbio.1000313. PubMed PMID: 20186266; PubMed Central PMCID: PMCPMC2826372.

31. Cascales E, Christie PJ. The versatile bacterial type IV secretion systems. Nat Rev Microbiol. 2003;1(2):137–49. doi: 10.1038/nrmicro753. PubMed PMID: 15035043; PubMed Central PMCID: PMCPMC3873781.

32. Costa TRD, Harb L, Khara P, Zeng L, Hu B, Christie PJ. Type IV secretion systems: Advances in structure, function, and activation. Mol Microbiol. 2021;115(3):436–52. Epub 20210107. doi: 10.1111/mmi.14670. PubMed PMID: 33326642; PubMed Central PMCID: PMCPMC8026593.

33. Wang J, Li J, Hou Y, Dai W, Xie R, Marquez-Lago TT, et al. BastionHub: a universal platform for integrating and analyzing substrates secreted by Gram-negative bacteria. Nucleic Acids Res. 2021;49(D1):D651–D9. doi: 10.1093/nar/gkaa899. PubMed PMID: 33084862; PubMed Central PMCID: PMCPMC7778982.

34. Patel V, Chevignon G, Manzano-Marin A, Brandt JW, Strand MR, Russell JA, et al. Cultivation-Assisted Genome of Candidatus Fukatsuia symbiotica; the Enigmatic “X-Type” Symbiont of Aphids. Genome Biol Evol. 2019;11(12):3510–22. Epub 2019/11/15. doi: 10.1093/gbe/evz252. PubMed PMID: 31725149; PubMed Central PMCID: PMCPMC7145644.

35. Patel V, Lynn-Bell N, Chevignon G, Kucuk RA, Higashi CHV, Carpenter M, et al. Mobile elements create strain-level variation in the services conferred by an aphid symbiont. Environ Microbiol. 2023;25(12):3333–48. Epub 20231020. doi: 10.1111/1462-2920.16520. PubMed PMID: 37864320.

36. Wu T, Rodrigues AA, Fayle TM, Henry LM. Defensive Symbiont Genotype Distributions Are Linked to Parasitoid Attack Networks. Ecol Lett. 2025;28(2):e70082. doi: 10.1111/ele.70082. PubMed PMID: 39964074; PubMed Central PMCID: PMCPMC11834374.

37. Granato ET, Meiller-Legrand TA, Foster KR. The Evolution and Ecology of Bacterial Warfare. Curr Biol. 2019;29(11):R521–R37. doi: 10.1016/j.cub.2019.04.024. PubMed PMID: 31163166.

38. Ballinger MJ, Perlman SJ. Generality of toxins in defensive symbiosis: Ribosome-inactivating proteins and defense against parasitic wasps in Drosophila. PLoS Pathog. 2017;13(7):e1006431. Epub 2017/07/07. doi: 10.1371/journal.ppat.1006431. PubMed PMID: 28683136; PubMed Central PMCID: PMCPMC5500355.

39. Goldstein EB, de Anda Acosta Y, Henry LM, Parker BJ. Variation in density, immune gene suppression, and coinfection outcomes among strains of the aphid endosymbiont Regiella insecticola. Evolution. 2023;77(7):1704–11. doi: 10.1093/evolut/qpad071. PubMed PMID: 37094805.

40. Chevignon G, Boyd BM, Brandt JW, Oliver KM, Strand MR. Culture-Facilitated Comparative Genomics of the Facultative Symbiont Hamiltonella defensa. Genome Biol Evol. 2018;10(3):786–802. Epub 2018/02/17. doi: 10.1093/gbe/evy036. PubMed PMID: 29452355; PubMed Central PMCID: PMCPMC5841374.

41. Monnin D, Jackson R, Kiers ET, Bunker M, Ellers J, Henry LM. Parallel Evolution in the Integration of a Coobligate Aphid Symbiosis. Curr Biol. 2020;30(10):1949–57 e6. Epub 2020/04/04. doi: 10.1016/j.cub.2020.03.011. PubMed PMID: 32243856.

42. Smee MR, Raines SA, Ferrari J. Genetic identity and genotype x genotype interactions between symbionts outweigh species level effects in an insect microbiome. ISME J. 2021;15(9):2537–46. Epub 20210312. doi: 10.1038/s41396-021-00943-9. PubMed PMID: 33712703; PubMed Central PMCID: PMCPMC8397793.

43. Foottit RG, Maw HE, CD Vond, Hebert PD. Species identification of aphids (Insecta: Hemiptera: Aphididae) through DNA barcodes. Mol Ecol Resour. 2008;8(6):1189–201. doi: 10.1111/j.1755-0998.2008.02297.x. PubMed PMID: 21586006.

44. Edgar RC. Muscle5: High-accuracy alignment ensembles enable unbiased assessments of sequence homology and phylogeny. Nat Commun. 2022;13(1):6968. Epub 20221115. doi: 10.1038/s41467-022-34630-w. PubMed PMID: 36379955; PubMed Central PMCID: PMCPMC9664440.

45. Camacho C, Coulouris G, Avagyan V, Ma N, Papadopoulos J, Bealer K, et al. BLAST+: architecture and applications. BMC Bioinformatics. 2009;10:421. Epub 20091215. doi: 10.1186/1471-2105-10-421. PubMed PMID: 20003500; PubMed Central PMCID: PMCPMC2803857.

46. Degnan PH, Moran NA. Evolutionary genetics of a defensive facultative symbiont of insects: exchange of toxin-encoding bacteriophage. Mol Ecol. 2008;17(3):916–29. Epub 2008/01/09. doi: 10.1111/j.1365-294X.2007.03616.x. PubMed PMID: 18179430.

47. Rausch T, Fritz MH, Untergasser A, Benes V. Tracy: basecalling, alignment, assembly and deconvolution of sanger chromatogram trace files. BMC Genomics. 2020;21(1):230. Epub 20200314. doi: 10.1186/s12864-020-6635-8. PubMed PMID: 32171249; PubMed Central PMCID: PMCPMC7071639.

48. Weldon SR, Strand MR, Oliver KM. Phage loss and the breakdown of a defensive symbiosis in aphids. Proc Biol Sci. 2013;280(1751):20122103. Epub 2012/11/30. doi: 10.1098/rspb.2012.2103. PubMed PMID: 23193123; PubMed Central PMCID: PMCPMC3574403.

49. Ewels P, Magnusson M, Lundin S, Kaller M. MultiQC: summarize analysis results for multiple tools and samples in a single report. Bioinformatics. 2016;32(19):3047–8. Epub 20160616. doi: 10.1093/bioinformatics/btw354. PubMed PMID: 27312411; PubMed Central PMCID: PMCPMC5039924.

50. Chen S, Zhou Y, Chen Y, Gu J. fastp: an ultra-fast all-in-one FASTQ preprocessor. Bioinformatics. 2018;34(17):i884–i90. doi: 10.1093/bioinformatics/bty560. PubMed PMID: 30423086; PubMed Central PMCID: PMCPMC6129281.

51. Kolmogorov M, Bickhart DM, Behsaz B, Gurevich A, Rayko M, Shin SB, et al. metaFlye: scalable long-read metagenome assembly using repeat graphs. Nat Methods. 2020;17(11):1103–10. Epub 20201005. doi: 10.1038/s41592-020-00971-x. PubMed PMID: 33020656; PubMed Central PMCID: PMCPMC10699202.

52. Prjibelski A, Antipov D, Meleshko D, Lapidus A, Korobeynikov A. Using SPAdes De Novo Assembler. Curr Protoc Bioinformatics. 2020;70(1):e102. doi: 10.1002/cpbi.102. PubMed PMID: 32559359.

53. Li H, Durbin R. Fast and accurate long-read alignment with Burrows-Wheeler transform. Bioinformatics. 2010;26(5):589–95. Epub 20100115. doi: 10.1093/bioinformatics/btp698. PubMed PMID: 20080505; PubMed Central PMCID: PMCPMC2828108.

54. Morgulis A, Coulouris G, Raytselis Y, Madden TL, Agarwala R, Schaffer AA. Database indexing for production MegaBLAST searches. Bioinformatics. 2008;24(16):1757–64. Epub 20080621. doi: 10.1093/bioinformatics/btn322. PubMed PMID: 18567917; PubMed Central PMCID: PMCPMC2696921.

55. Buchfink B, Reuter K, Drost HG. Sensitive protein alignments at tree-of-life scale using DIAMOND. Nat Methods. 2021;18(4):366–8. Epub 20210407. doi: 10.1038/s41592-021-01101-x. PubMed PMID: 33828273; PubMed Central PMCID: PMCPMC8026399.

56. Laetsch DR, Blaxter ML. BlobTools: Interrogation of genome assemblies. F1000Research. 2017;6. doi: 10.12688/f1000research.12232.1.

57. Li H, Handsaker B, Wysoker A, Fennell T, Ruan J, Homer N, et al. The Sequence Alignment/Map format and SAMtools. Bioinformatics. 2009;25(16):2078–9. Epub 2009/06/10. doi: 10.1093/bioinformatics/btp352. PubMed PMID: 19505943; PubMed Central PMCID: PMCPMC2723002.

58. Wick RR, Schultz MB, Zobel J, Holt KE. Bandage: interactive visualization of de novo genome assemblies. Bioinformatics. 2015;31(20):3350–2. Epub 20150622. doi: 10.1093/bioinformatics/btv383. PubMed PMID: 26099265; PubMed Central PMCID: PMCPMC4595904.

59. Parks DH, Imelfort M, Skennerton CT, Hugenholtz P, Tyson GW. CheckM: assessing the quality of microbial genomes recovered from isolates, single cells, and metagenomes. Genome Res. 2015;25(7):1043–55. Epub 20150514. doi: 10.1101/gr.186072.114. PubMed PMID: 25977477; PubMed Central PMCID: PMCPMC4484387.

60. Eddy SR. Accelerated Profile HMM Searches. PLoS Comput Biol. 2011;7(10):e1002195. Epub 20111020. doi: 10.1371/journal.pcbi.1002195. PubMed PMID: 22039361; PubMed Central PMCID: PMCPMC3197634.

61. Guindon S, Dufayard JF, Lefort V, Anisimova M, Hordijk W, Gascuel O. New algorithms and methods to estimate maximum-likelihood phylogenies: assessing the performance of PhyML 3.0. Syst Biol. 2010;59(3):307–21. Epub 20100329. doi: 10.1093/sysbio/syq010. PubMed PMID: 20525638.

62. Letunic I, Bork P. Interactive Tree Of Life (iTOL) v5: an online tool for phylogenetic tree display and annotation. Nucleic Acids Res. 2021;49(W1):W293–W6. doi: 10.1093/nar/gkab301. PubMed PMID: 33885785; PubMed Central PMCID: PMCPMC8265157.

63. Lee MD. GToTree: a user-friendly workflow for phylogenomics. Bioinformatics. 2019;35(20):4162–4. doi: 10.1093/bioinformatics/btz188. PubMed PMID: 30865266; PubMed Central PMCID: PMCPMC6792077.

64. Capella-Gutierrez S, Silla-Martinez JM, Gabaldon T. trimAl: a tool for automated alignment trimming in large-scale phylogenetic analyses. Bioinformatics. 2009;25(15):1972–3. Epub 20090608. doi: 10.1093/bioinformatics/btp348. PubMed PMID: 19505945; PubMed Central PMCID: PMCPMC2712344.

65. Hyatt D, Chen GL, Locascio PF, Land ML, Larimer FW, Hauser LJ. Prodigal: prokaryotic gene recognition and translation initiation site identification. BMC Bioinformatics. 2010;11:119. Epub 20100308. doi: 10.1186/1471-2105-11-119. PubMed PMID: 20211023; PubMed Central PMCID: PMCPMC2848648.

66. Kalyaanamoorthy S, Minh BQ, Wong TKF, von Haeseler A, Jermiin LS. ModelFinder: fast model selection for accurate phylogenetic estimates. Nat Methods. 2017;14(6):587–9. Epub 20170508. doi: 10.1038/nmeth.4285. PubMed PMID: 28481363; PubMed Central PMCID: PMCPMC5453245.

67. Hoang DT, Chernomor O, von Haeseler A, Minh BQ, Vinh LS. UFBoot2: Improving the Ultrafast Bootstrap Approximation. Mol Biol Evol. 2018;35(2):518–22. doi: 10.1093/molbev/msx281. PubMed PMID: 29077904; PubMed Central PMCID: PMCPMC5850222.

68. Tange O. GNU Parallel 2018.

69. Nguyen LT, Schmidt HA, von Haeseler A, Minh BQ. IQ-TREE: a fast and effective stochastic algorithm for estimating maximum-likelihood phylogenies. Mol Biol Evol. 2015;32(1):268–74. Epub 20141103. doi: 10.1093/molbev/msu300. PubMed PMID: 25371430; PubMed Central PMCID: PMCPMC4271533.

70. Stamatakis A. RAxML version 8: a tool for phylogenetic analysis and post-analysis of large phylogenies. Bioinformatics. 2014;30(9):1312–3. Epub 20140121. doi: 10.1093/bioinformatics/btu033. PubMed PMID: 24451623; PubMed Central PMCID: PMCPMC3998144.

71. Seemann T. Prokka: rapid prokaryotic genome annotation. Bioinformatics. 2014;30(14):2068–9. Epub 20140318. doi: 10.1093/bioinformatics/btu153. PubMed PMID: 24642063.

72. Page AJ, Cummins CA, Hunt M, Wong VK, Reuter S, Holden MT, et al. Roary: rapid large-scale prokaryote pan genome analysis. Bioinformatics. 2015;31(22):3691–3. Epub 20150720. doi: 10.1093/bioinformatics/btv421. PubMed PMID: 26198102; PubMed Central PMCID: PMCPMC4817141.

73. Hadfield J, Croucher NJ, Goater RJ, Abudahab K, Aanensen DM, Harris SR. Phandango: an interactive viewer for bacterial population genomics. Bioinformatics. 2018;34(2):292–3. doi: 10.1093/bioinformatics/btx610. PubMed PMID: 29028899; PubMed Central PMCID: PMCPMC5860215.

74. Hernandez-Plaza A, Szklarczyk D, Botas J, Cantalapiedra CP, Giner-Lamia J, Mende DR, et al. eggNOG 6.0: enabling comparative genomics across 12 535 organisms. Nucleic Acids Res. 2023;51(D1):D389–D94. doi: 10.1093/nar/gkac1022. PubMed PMID: 36399505; PubMed Central PMCID: PMCPMC9825578.

75. Arndt D, Grant JR, Marcu A, Sajed T, Pon A, Liang Y, et al. PHASTER: a better, faster version of the PHAST phage search tool. Nucleic Acids Res. 2016;44(W1):W16–21. Epub 20160503. doi: 10.1093/nar/gkw387. PubMed PMID: 27141966; PubMed Central PMCID: PMCPMC4987931.

76. Wang RH, Yang S, Liu Z, Zhang Y, Wang X, Xu Z, et al. PhageScope: a well-annotated bacteriophage database with automatic analyses and visualizations. Nucleic Acids Res. 2024;52(D1):D756–D61. doi: 10.1093/nar/gkad979. PubMed PMID: 37904614; PubMed Central PMCID: PMCPMC10767790.

77. Baril T, Galbraith J, Hayward A. Earl Grey: A Fully Automated User-Friendly Transposable Element Annotation and Analysis Pipeline. Mol Biol Evol. 2024;41(4). doi: 10.1093/molbev/msae068. PubMed PMID: 38577785; PubMed Central PMCID: PMCPMC11003543.

78. Aramaki T, Blanc-Mathieu R, Endo H, Ohkubo K, Kanehisa M, Goto S, et al. KofamKOALA: KEGG Ortholog assignment based on profile HMM and adaptive score threshold. Bioinformatics. 2020;36(7):2251–2. doi: 10.1093/bioinformatics/btz859. PubMed PMID: 31742321; PubMed Central PMCID: PMCPMC7141845.

79. Kanehisa M, Sato Y. KEGG Mapper for inferring cellular functions from protein sequences. Protein Sci. 2020;29(1):28–35. Epub 20190829. doi: 10.1002/pro.3711. PubMed PMID: 31423653; PubMed Central PMCID: PMCPMC6933857.

80. Krzywinski M, Schein J, Birol I, Connors J, Gascoyne R, Horsman D, et al. Circos: an information aesthetic for comparative genomics. Genome Res. 2009;19(9):1639–45. Epub 20090618. doi: 10.1101/gr.092759.109. PubMed PMID: 19541911; PubMed Central PMCID: PMCPMC2752132.

81. Parker BJ, McLean AHC, Hrcek J, Gerardo NM, Godfray HCJ. Establishment and maintenance of aphid endosymbionts after horizontal transfer is dependent on host genotype. Biol Lett. 2017;13(5). Epub 2017/06/02. doi: 10.1098/rsbl.2017.0016. PubMed PMID: 28566541; PubMed Central PMCID: PMCPMC5454236.

82. Parker BJ, Hrcek J, McLean AHC, Godfray HCJ. Genotype specificity among hosts, pathogens, and beneficial microbes influences the strength of symbiont-mediated protection. Evolution. 2017;71(5):1222–31. Epub 2017/03/03. doi: 10.1111/evo.13216. PubMed PMID: 28252804; PubMed Central PMCID: PMCPMC5516205.

83. Parker BJ, Garcia JR, Gerardo NM. Genetic variation in resistance and fecundity tolerance in a natural host-pathogen interaction. Evolution. 2014;68(8):2421–9. Epub 2014/04/03. doi: 10.1111/evo.12418. PubMed PMID: 24689981.

84. Kolp MR, de Anda Acosta Y, Brewer W, Nichols HL, Goldstein EB, Tallapragada K, et al. Pathogenmicrobiome interactions and the virulence of an entomopathogenic fungus. Appl Environ Microbiol. 2024:e0229323. Epub 20240524. doi: 10.1128/aem.02293-23. PubMed PMID: 38786361.

85. Bender W, Spierer P, Hogness DS. Chromosomal Walking and Jumping to Isolate DNA from the Ace and rosy Loci and the Bithorax Complex in Drosophila melanogaster. J Mol Biol. 1983;168:17–33.

86. Team RC. R: A language and environment for statistical computing.: R Foundation for Statistical Computing; 2017.

87. Parker BJ, Barribeau SM, Laughton AM, Griffin LH, Gerardo NM. Life-history strategy determines constraints on immune function. J Anim Ecol. 2017;86(3):473–83. Epub 2017/02/18. doi: 10.1111/1365-2656.12657. PubMed PMID: 28211052.

88. Anders S, Pyl PT, Huber W. HTSeq--a Python framework to work with high-throughput sequencing data. Bioinformatics. 2015;31(2):166–9. Epub 2014/09/28. doi: 10.1093/bioinformatics/btu638. PubMed PMID: 25260700; PubMed Central PMCID: PMCPMC4287950.

89. Robinson MD, McCarthy DJ, Smyth GK. edgeR: a Bioconductor package for differential expression analysis of digital gene expression data. Bioinformatics. 2010;26(1):139–40. Epub 2009/11/17. doi: 10.1093/bioinformatics/btp616. PubMed PMID: 19910308; PubMed Central PMCID: PMCPMC2796818.

