## Supplementary figures and images for "Phenotypic divergence is driven by mobile genetic elements in a heritable insect symbiont"

### Figure S1

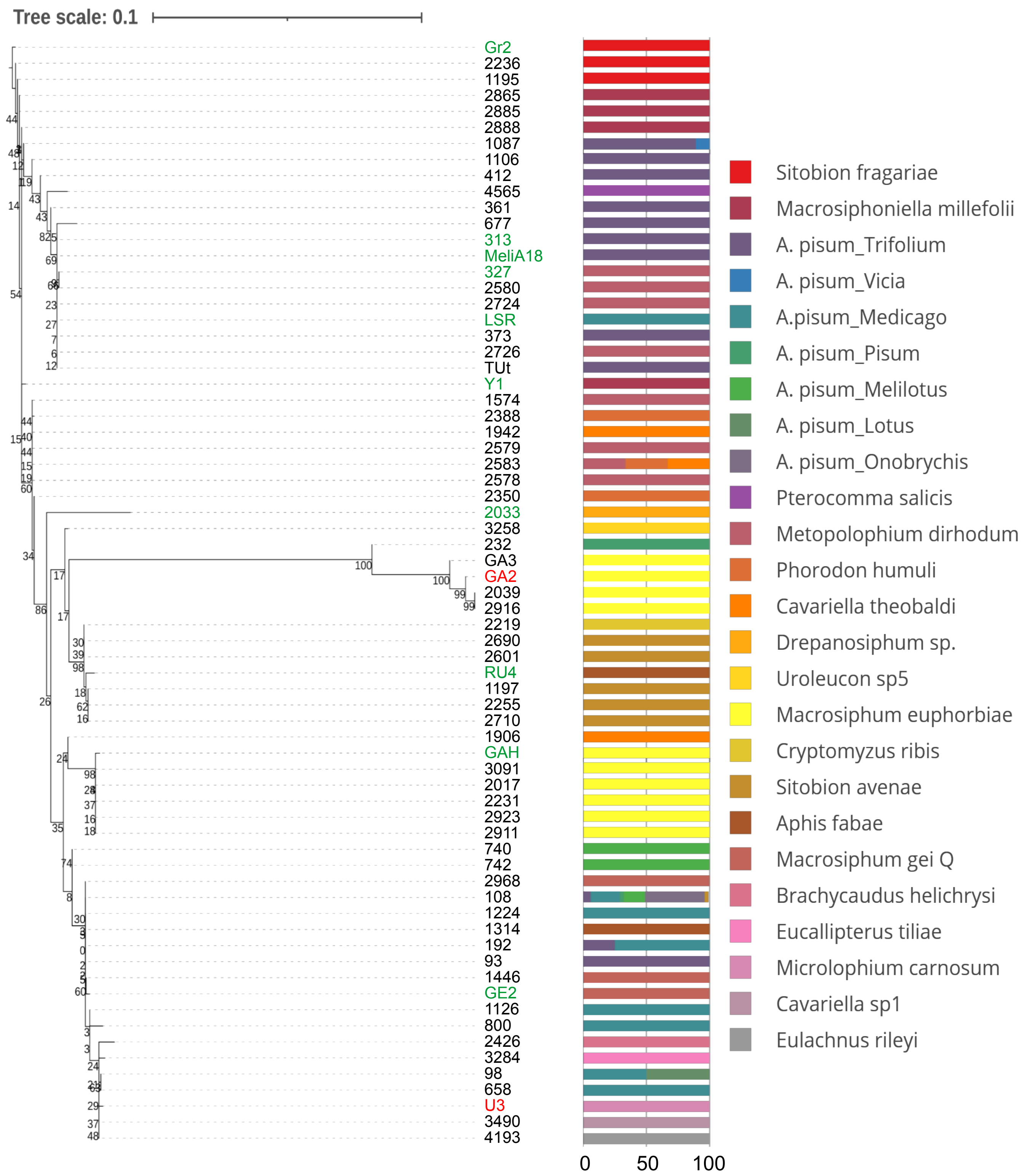

### Figure S2

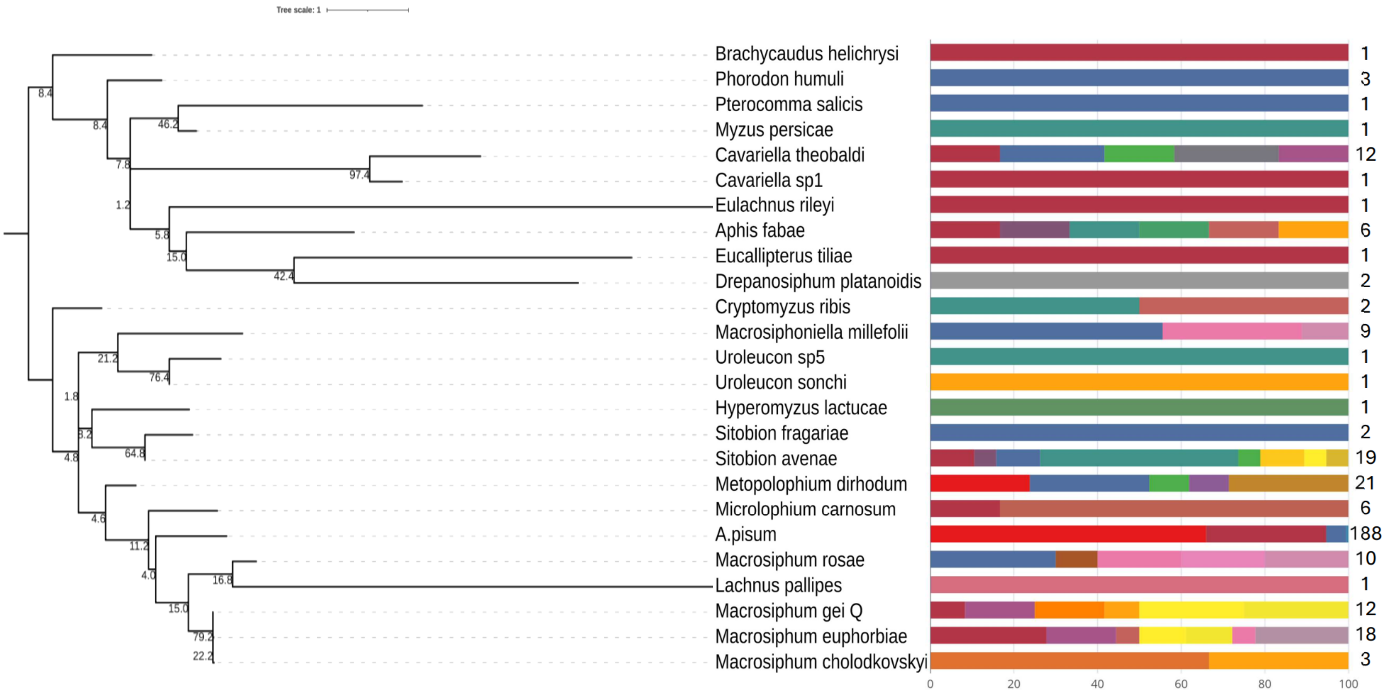

### Figure S3

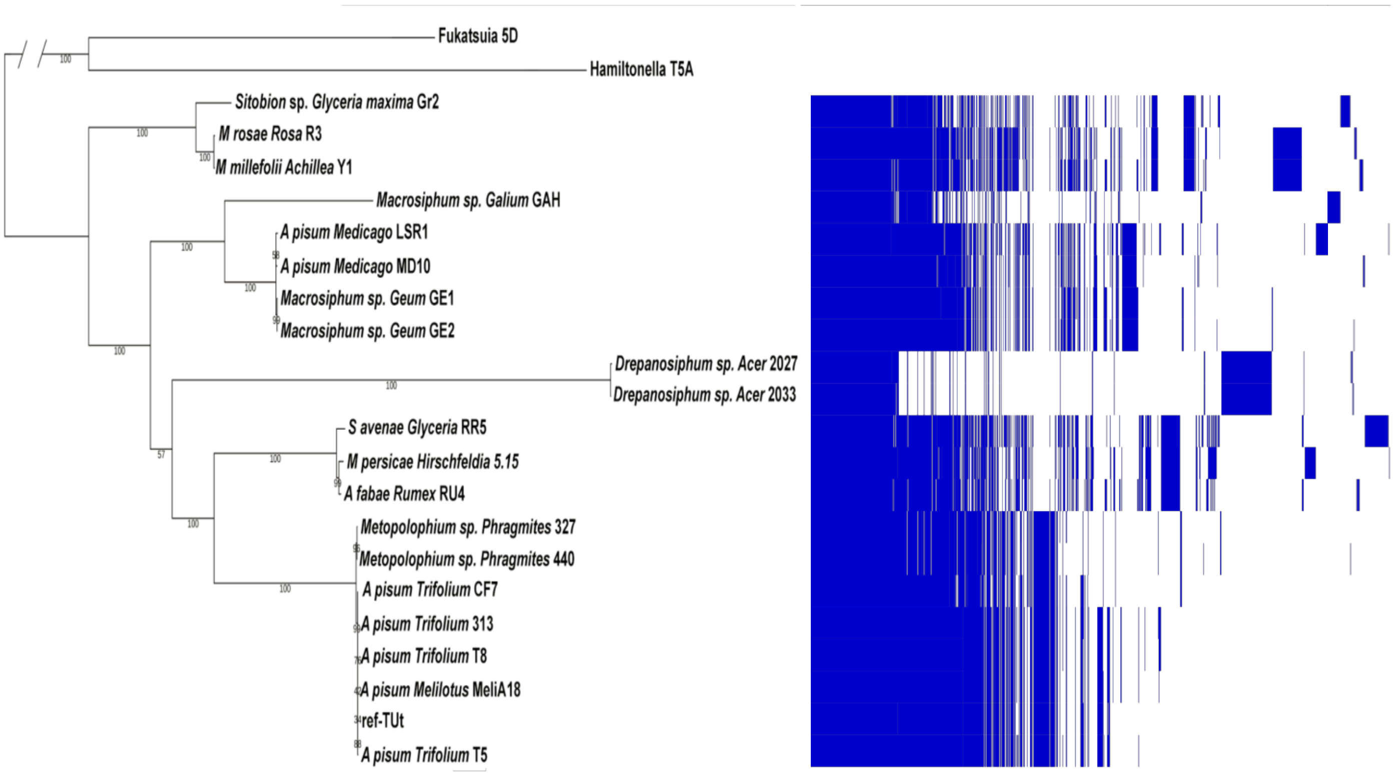

### Figure S4

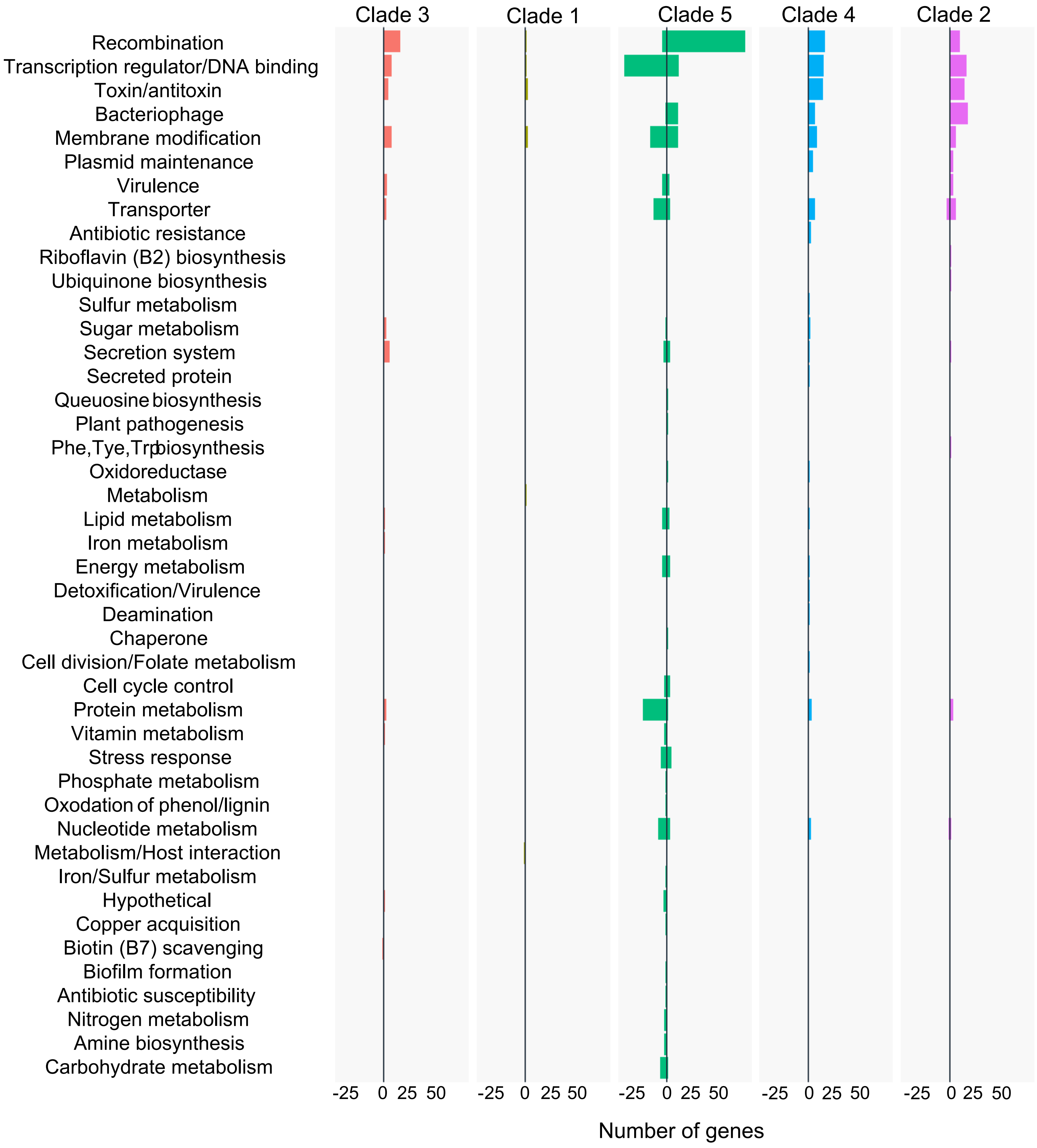

### Figure S5

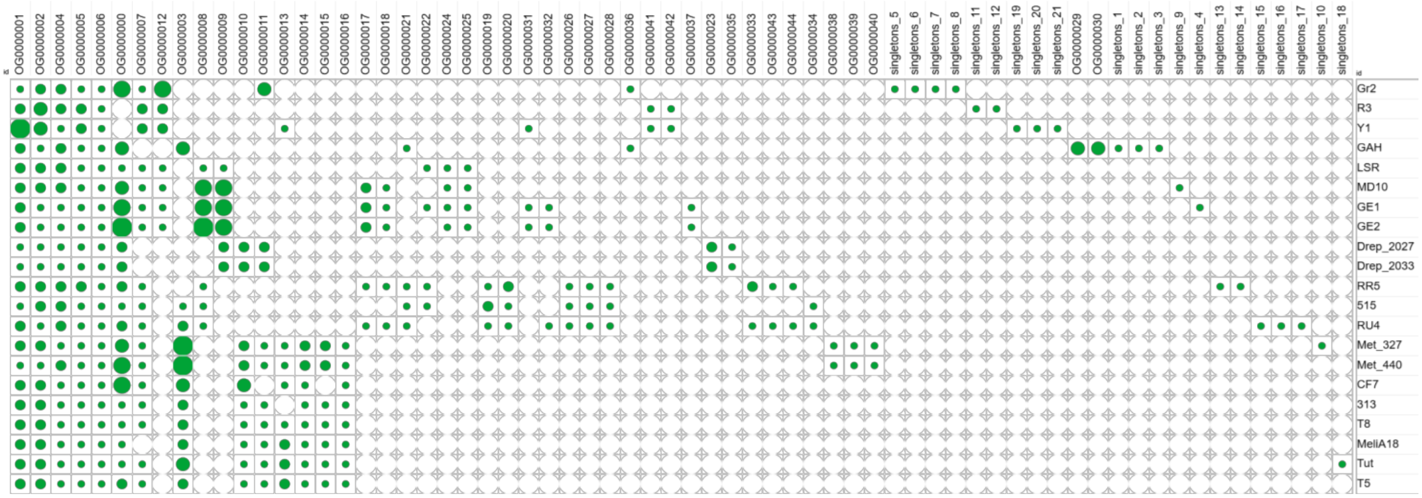

### Figure S6

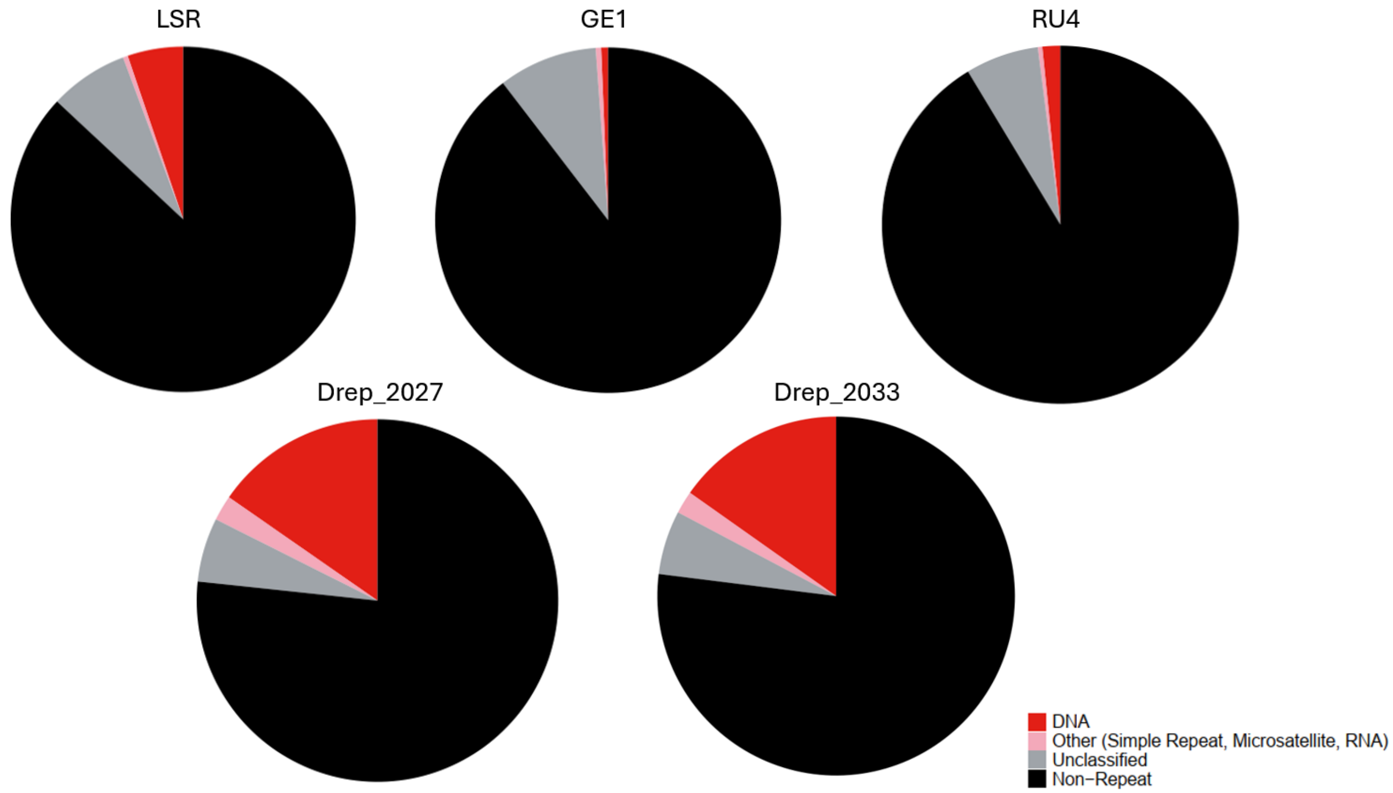

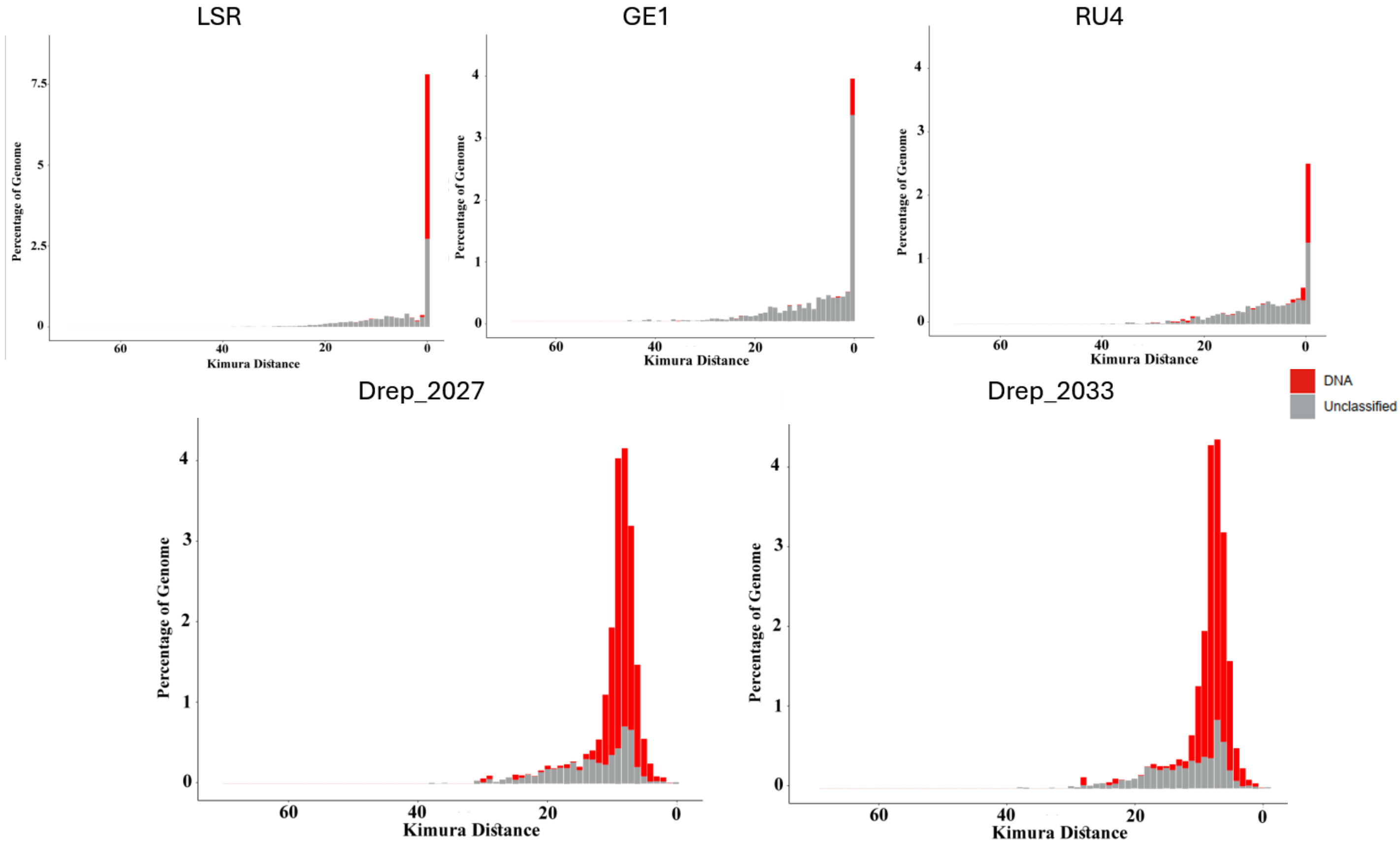

### Figure S7

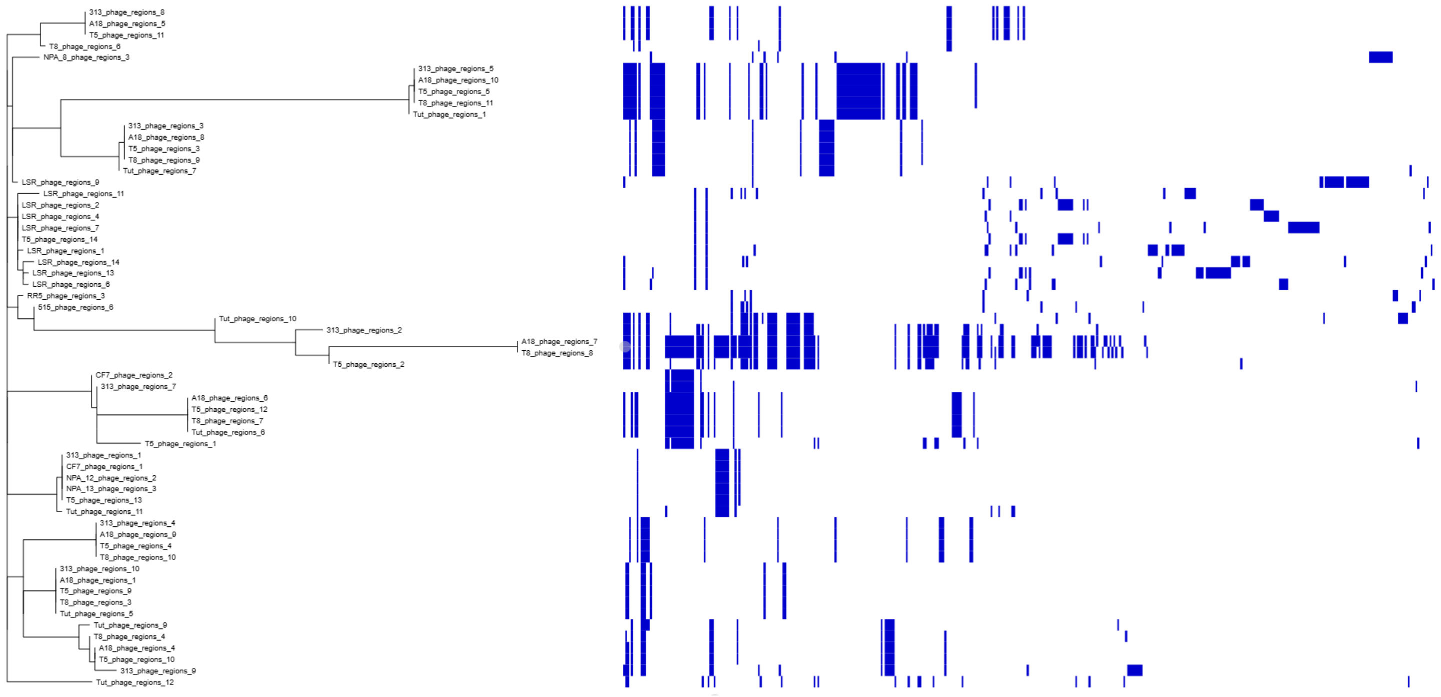

### Figure S8

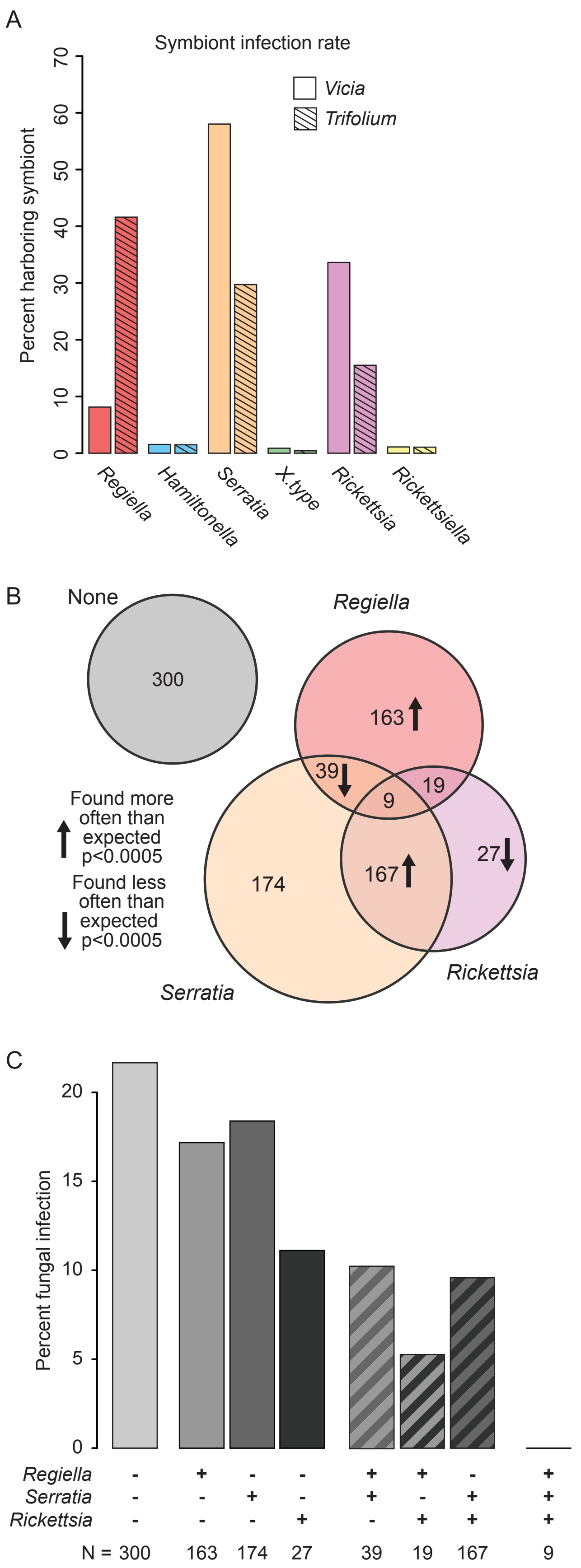

### Figure S9

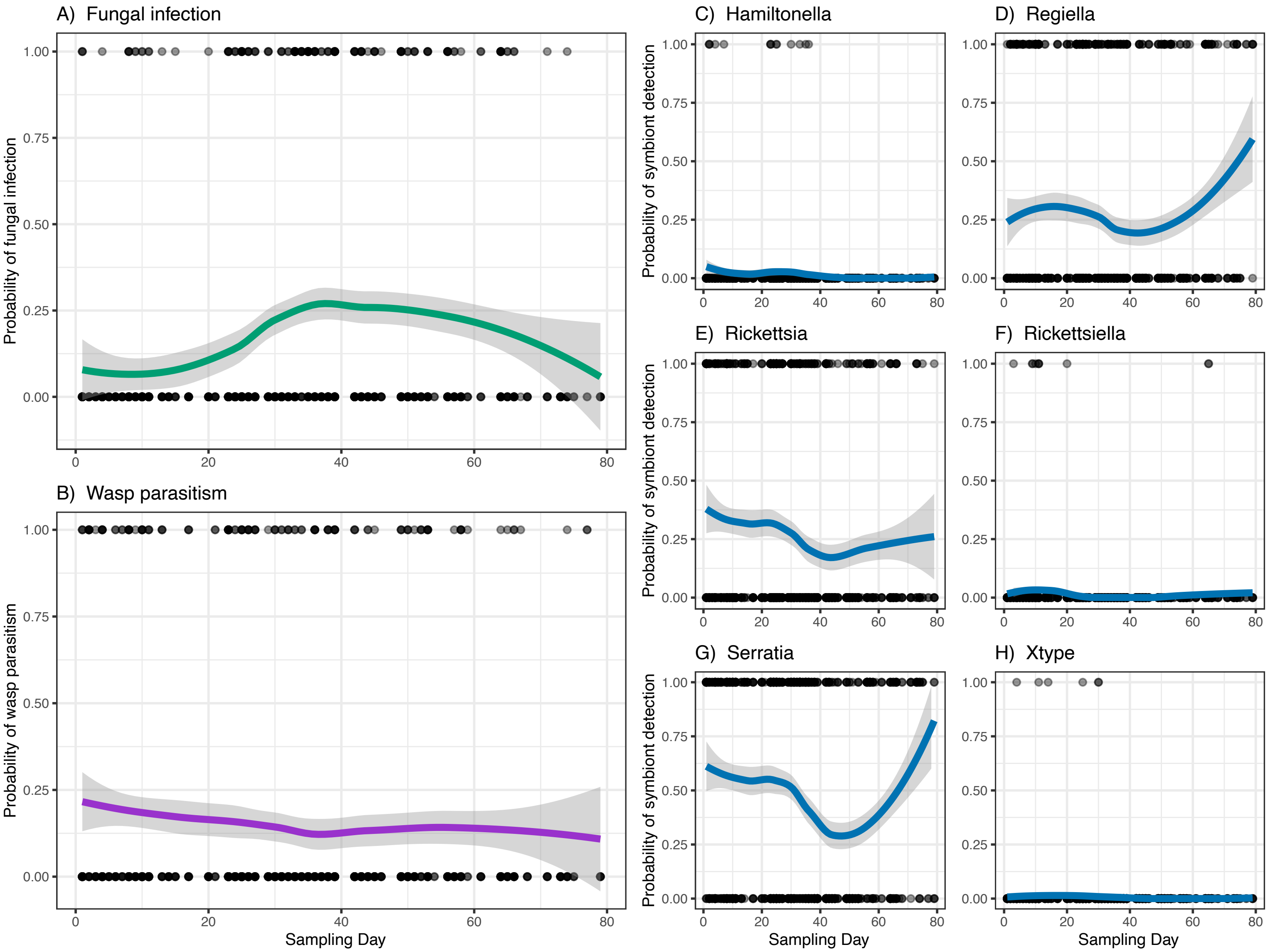
